# Effect of Azithromycin treatment on the microbial composition, functional dynamics and resistomes of endocervical, vaginal and rectal microbiomes of women in Fiji with *Chlamydia trachomatis* infection

**DOI:** 10.1101/2025.04.02.646699

**Authors:** Sankhya Bommana, Olusola Olagoke, Yi-Juan Hu, Roselle Wang, Mike Kama, Morgan Dehdashti, Reshma Kodimerla, Timothy D. Read, Deborah Dean

## Abstract

Antibiotics disrupt mucosal microbial communities, yet the effects on microbiomes with *Chlamydia trachomatis* (*Ct*) infection remain poorly understood. Some data exist on vaginal microbiomes pre- and post-treatment, but none are available for the endocervix or rectum that are primary sites of infection. We applied metagenomic shotgun sequencing to vaginal, endocervical and rectal samples from women who, overtime, had *Ct* persistence, clearance, or no infection to evaluate azithromycin-induced changes in microbial composition, function, and the resistome. Our results show a shift in composition and function that support *Ct* post-treatment with azithromycin resistance mutations in the *Ct rpl*V gene and significant endocervical enrichment of azithromycin resistance genes in *Lactobacillus iners* and *Gardnerella vaginalis*, the strains of which have moderate/high potential for biofilm formation. These findings highlight the unintended ecological consequences of azithromycin, including resistance gene propagation and microbiome shifts that support persistent/recurrent *Ct*, emphasizing the need for novel treatment and microbiome- preserving strategies.

## Introduction

*Chlamydia trachomatis* (*Ct*) is the most prevalent bacterial sexually transmitted infection worldwide^1^. Pacific Island countries and territories (PICT) of the Western Pacific Region have some of the highest reported global rates of *Ct* sexually transmitted infections (STIs)^2,3^. In Fiji, the most recent data from 2023 showed a prevalence of 29.4% among women aged 18 to 24 years and 17% for those 25 to 40 years^4^. This high prevalence is due, in part, to syndromic management, which relies on patient symptoms and easily identifiable clinical signs to determine the need for antibiotic treatment^5^. However, this approach is especially problematic for STIs such as *Ct* where up to 80% of the cases are asymptomatic^6–8^. Indeed, syndromic management has missed approximately 90% of *Ct* infections in Fiji^8^, 77.8% in South Africa^9^ and 80% in Papua New Guinea^10^. As a result, untreated *Ct* infections can progress to upper genital tract sequelae, including pelvic inflammatory disease, tubal factor infertility, ectopic pregnancy and preterm birth^2,3,11–14^ along with unchecked transmission to partners.

A healthy vaginal microbiome is crucial for preventing STIs and is classified based on five community state types (CSTs) and 15 subCSTs^15–18^. subCST I-A and I-B, CST II, CST III-A and III-B and CST-V are dominated by *Lactobacillus crispatus, L. gasseri, L. iners,* and *L. jensenii*, respectively. In general, *Lactobacillus* species produce lactic acid, bacteriocins, and other antimicrobial compounds that restrict infection by most pathogenic bacteria^15–18^. CST IVs are abundant in diverse facultative and strict anaerobic organisms such as *Gardnerella, Atopobium, Megasphaera, Prevotella,* and *Mobiluncus* with a dearth of *Lactobacillus* spp. and include five previous and four new subCSTs. The latter four were identified using metagenomic shotgun sequences (MSS) and the relative abundance of bacteria from the microbiomes of diverse Pacific Islander ethnicities in Fiji^16^. Because of the lack of protective effects of most *Lactobacillus* spp., except for *L. iners*, subCST IIIs and subCST IVs are predisposed to bacterial vaginosis (BV) with an increased risk of *Ct* and other STIs like *Neisseria gonorrhoeae (Ng)*, herpes simplex virus type 2 (HSV-2), human Papillomavirus (HPV), and human immunodeficiency virus (HIV-1)^19–21^.

A few studies have shown that the vaginal microbiota responds to antimicrobial treatment with shifts in microbial composition^22–24^. Using 16S rRNA sequencing of vaginal samples from Asian women, one study reported that *Ct* infections treated with azithromycin can lead to an *L. iners*-dominant microbiome (i.e., subCST III-A or III-B) that transitioned from a higher relative abundance of *Enterobacter, Prevotella,* and *Streptococcus* spp. associated with subCST IVs^24^. This post-treatment abundance of *L. iners* was postulated to be a result of *L. iners* resistance to azithromycin. Unlike other *Lactobacillus* species, *L. iners* is unable to produce D (-) lactic acid, but can downregulate histone deacetylase 4, which typically prevents *Ct* induced cell proliferation^25^. As a result, *L. iners* is predicted to create a vaginal microenvironment susceptible to *Ct* reinfection and to other STIs^25,26^. This is consistent with another study that showed an association between *L. iners*-dominated microbiomes and incident *Ct* infection^19^.

Given the limited data on the effects of treatment on vaginal microbiomes that, to date, have relied on 16S rRNA sequencing, and none on the endocervical or rectal microbiomes, we employed MSS to investigate the impact of azithromycin on the microbial composition, functional dynamics and resistomes of paired vaginal, endocervical and rectal microbiomes among *Ct*-infected women in Fiji who were treated at baseline and then seen for post-treatment follow-up. These data were compared to uninfected, untreated women in any site at either time point.

## Results

### Patient and sample characteristics, and metagenomic shotgun sequencing data

Women 18 to 40 years of age attending Fiji Ministry of Health and Medical Services (MHMS) health centers and two university clinics were enrolled in the parent study as described^4,27^. Three cohorts with paired endocervical, vaginal and rectal samples were created based on presence/absence of *Ct* infection and their clearance or persistence of infection three months following azithromycin treatment, including one cohort of women who remained free of infection and were not treated (**Figure 1**).

**Figure 1.**
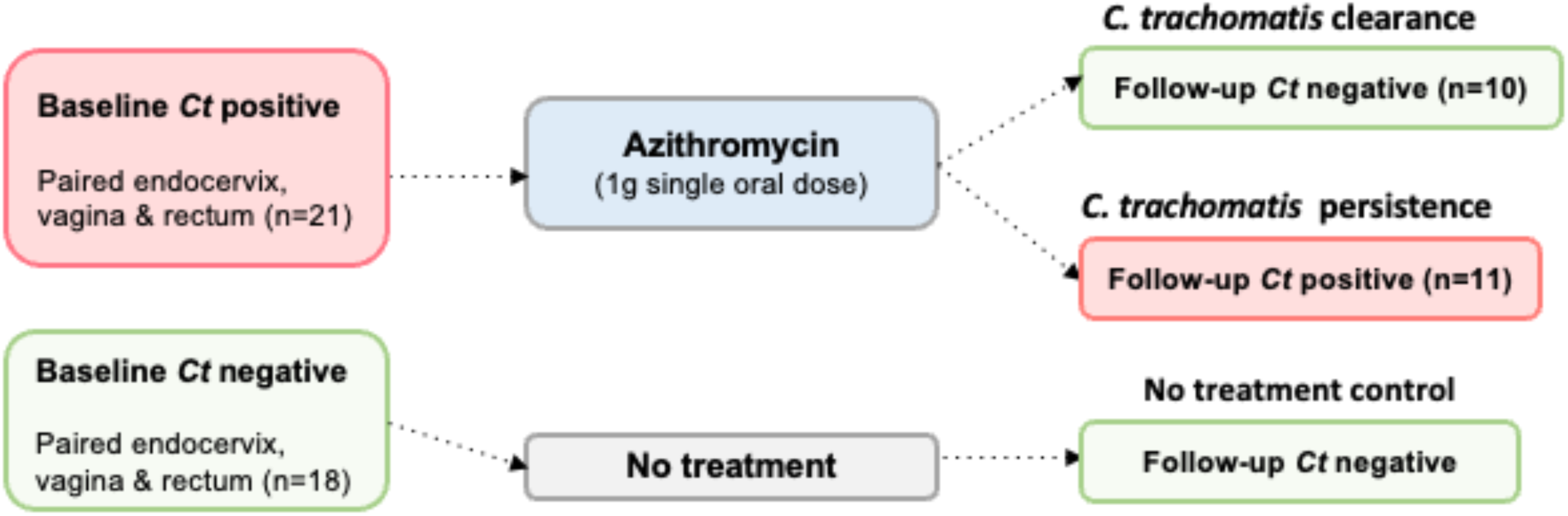
Schematic representation of the three cohorts of Pacific Islander women residing in Fiji, *Ct* status at baseline and follow-up at 3 months, anatomic sites tested, and treatment regime. *Ct*, *C. trachomatis*.

The vaginal, endocervical and rectal metagenome data were pre-processed as previously described^16,28^ and yielded a total of 7.3 billion, 7.9 billion and 3.4 billion raw reads, respectively, of which 6.4 billion (87.19%), 7.2 billion (91.42%) and 1.2 billion (35.92%) were identified as human contamination, respectively (**Table S1**).

The metadata for the study are shown in **Table S2** and were provided by the parent study^4,27^ including infection status for *Ct* and *Ng* based on the Xpert CT/NG assay. *Ng*, *Mycoplasma genitalium* (*Mg*), *Trichomonas vaginalis* (*Tv*) and low risk (lr) and high risk (hr) Human Papilloma Virus (HPV) types were identified from the MSS data (see Methods).

For the *Ct* persistence cohort, *Ng* cases increased from 36.3% to 45.5% for vaginal, 63.6% to 81.8% for endocervical and 36.3% to 63.6% for rectal microbiomes at baseline and follow-up, respectively (**Table S2**). The proportion of endocervical *Ng* cases in this cohort at follow-up were significantly higher than the untreated cohort (*P*<0.021; OR: 9; 95% CI: 1.46-55.5) (**Figure 2A**). In the *Ct* clearance cohort, vaginal *Ng* cases were like the persistence cohort (40%), but lower for the endocervix (30%) at both baseline and follow-up with no rectal *Ng* infections. Among controls, *Ng* cases were low and similar at baseline and follow-up with no rectal *Ng* infections at follow up. Very few *Mg* and *Tv* cases were detected in any cohort (**Table S2**).

**Figure 2.**
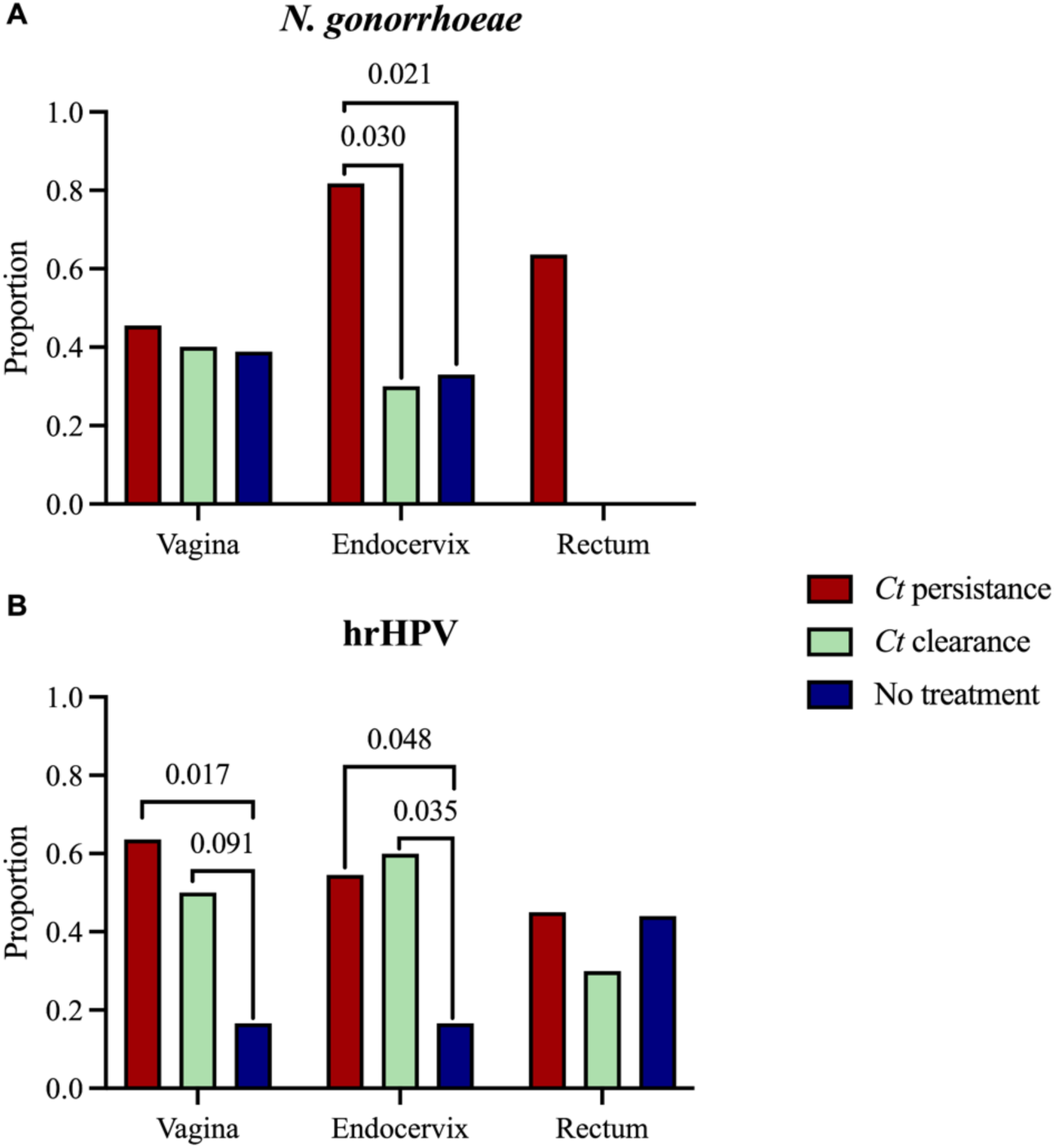
Proportion of samples with N. *gonorrhoeae* (A) or high risk (hr)HPV (B) at follow-up compared between treatment cohort and the untreated, uninfected control cohort. *P* values are shown when <0.1. There were no *N. gonorrhoeae* infections in the rectum at follow-up for the *Ct* clearance and no treatment cohorts.

The distribution of lrHPV and hrHPV in the paired vaginal, endocervical and rectal microbiomes for the three cohort datasets are shown in **Table S3**. There were no significant differences at baseline or at follow-up in the proportion of hrHPV types between the three anatomic sites or between baseline and follow-up for the respective cohort. While women at baseline were not significantly more likely to have the same hrHPV type at follow up, women in both the *Ct* persistent and *Ct* clearance cohorts were significantly more likely to have any hrHPV type in the endocervix (*P*<0.048; OR: 6.1 ; 95% CI: 1.1-33.3; and P<0.035; OR7.5; 95% CI: 1.28-44.1, respectively) and, for the *Ct* persistence cohort, in the vagina (*P*<0.017; OR: 8.6; 95% CI: 1.5-50.1) following treatment compared to uninfected women who were not treated (**Figure 2B**).

### Emergence of pathobionts in the endocervical and rectal microbiomes following azithromycin treatment contribute to Ct persistence

The Linear Decomposition Model (LDM)^29^ was employed to identify species associated with *Ct* clearance and persistence in comparison to the no treatment control cohort at baseline and at follow-up in the three paired anatomic sites (**Figure 3**). While there were no differences at baseline, several species were significantly associated with *Ct* persistence at follow up, including *Ng* in the endocervix, *Leptotrichia hofstadil* in the vagina, and several BV-associated and other species in the rectum such as *Ng, Streptococcus lutetiensis*, *Prevotella* spp. CAG 1092 and CAG 5226, *Ruminococcus* sp*. CAG 330, Prevotella* sp. 885, *Actinomyces naeslundii* and *Fusobacterium mortiferum* (**Figure 3**).

**Figure 3.**
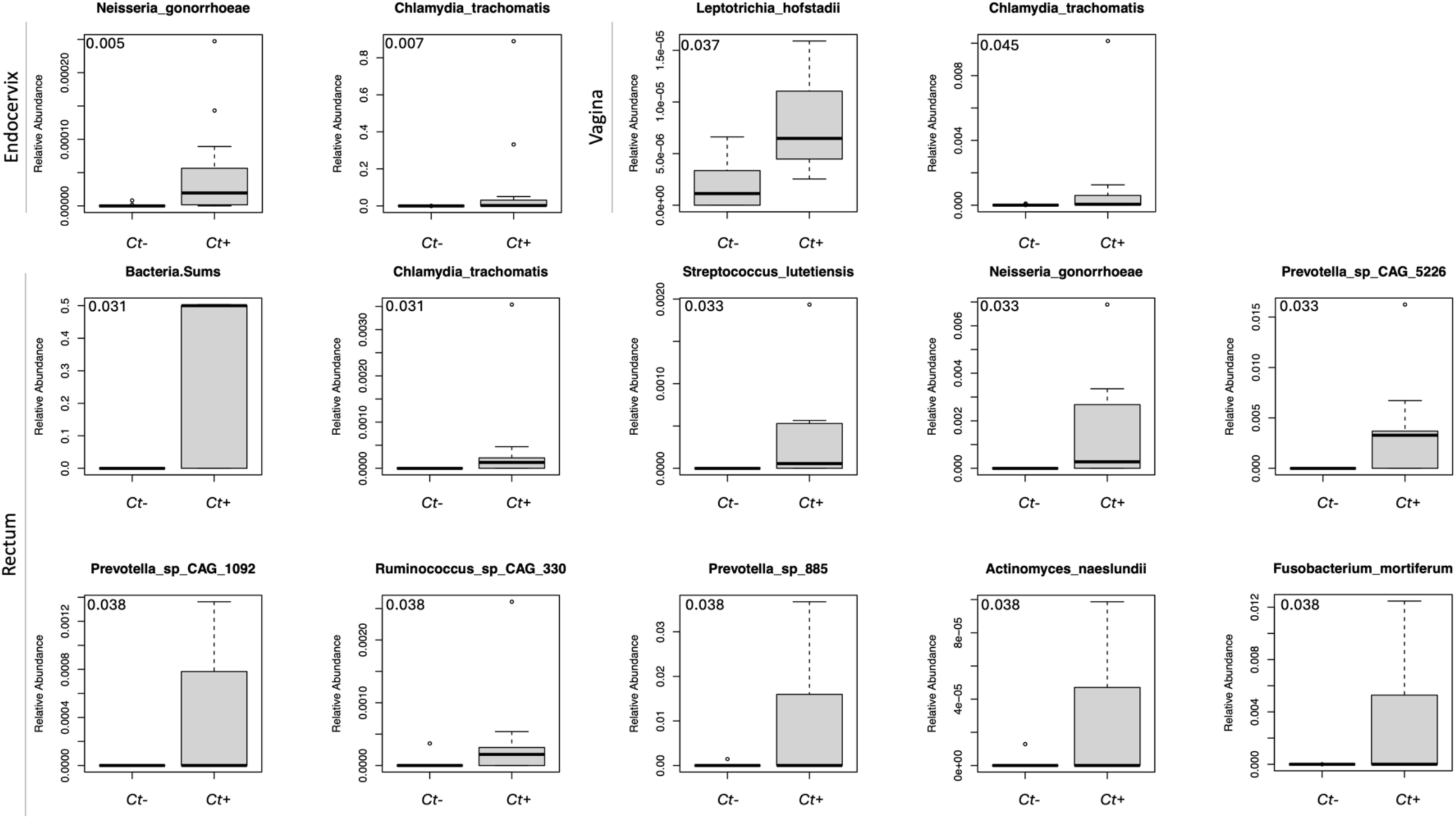
LDM analysis of significant differential species in the *Ct* persistence cohort datasets compared to the no treatment control cohort at follow-up for the paired endocervical, rectal and vaginal microbiomes. No significant differences were found at the baseline timepoint. Only species with a significant adjusted *P* value <0.05 (upper left corner of each bar graph) are represented.

### Species predictive of persistent C. trachomatis infection following azithromycin treatment

We also employed the LDM^29^ to identify taxa that changed in relative abundance before and after azithromycin treatment. While there are no published biomarkers at baseline that would predict clearance or persistent *Ct* infection following treatment, we found that *Campylobacter hominis* (*P*=0.005), *Coprococcus catus* (*P*=0.005), *Ruminococcus callidus* (*P*=0.0471) and *Roseburia inulinivorans* (*P*=0.0471), and a decrease in *Lactobacillus acidophilus* (*P*=0.0075) and

*Lactobacillus fermentum* (*P*=0.012), were significant baseline predictors for women who developed *Ct* persistence at follow up, but only in the rectum. *Ruminococcus* spp. are associated with gut microbiome enterotype 3 while *L. acidophilus* and *L. fermentum* are inversely correlated with enterotype 1^30^. The baseline rectal microbial diversity for the *Ct* persistence cohort was significantly different from the *Ct* clearance (*P*=0.002) and control cohorts (*P*=0.0002).

In comparing the relative abundance of the species in each anatomic site, there was a significant effect of time between baseline and follow-up, and a significant difference in the microbiome transition in each anatomic site based on the interaction of time and anatomic site (*P*=0.0002). The species responsible for these differences were *Acidaminococcus fermentans*, *Agathobaculum butyriciproducens, Anaerococcus vaginalis* and several species from the genus *Actinomyces* (all at *P*=3.24e-05).

Overall, the cohort datasets of paired vaginal, endocervical and rectal sites did not show any significant difference in alpha diversity between baseline and follow up for any cohort, although there was a decrease in diversity for the *Ct* clearance cohort (**Figure S1**). Among women whose *Ct* infection persisted, there was a slight increase in species richness and Shannon index from baseline to follow-up, particularly in the vagina and rectum (**Figure S1**). There was also no significant difference in beta diversity between baseline and follow up for any cohort (**Figure S2**).

### Comparison of subCST and mgCST classification systems revealed lack of well-defined and characterized mgCST categories for Pacific Islander microbiomes

The overall accuracy of the metagenomic (mg)CST system^31^ was compared with the expanded subCST system that we developed in 2023 for classification and functional associations of the Pacific Islander endocervical and vaginal microbiomes^16^. The microbiomes of the three cohort datasets that included paired endocervical and vaginal samples (**Figure 1**) and another cohort with only vaginal samples (see Methods) represented mainly subCST III-A and III-B, and IV-B, IV-D0, IV-D1, IV-D2 and IV-E. Common mgCSTs were 10 (mgS: *L. iners* 1), 12 (mgS: *L. iners* 3), 20 (mgS: *Gardnerella vaginalis* 1) and 24 (mgS: *G. vaginalis* 4) (**Table S4**).

Among Pacific Islander microbiomes, 10.5% (22/209) of vaginal and 25.6% (20/78) of endocervical microbiomes had mgCST scores of less than 0.5, indicating low similarity to the reference centroid of the assigned mgCST (**Table S4**). In the Holm *et al.* study^31^, the lowest similarity scores (<0.5) were attributed mainly to mgCST 27, which represents ‘other’ bacteria that are not defined. The assigned mgCSTs with low scores among vaginal microbiomes were 6, 25, and 27; the latter had 40.9% (9/22) of the low scores (**Figure 4**). In the endocervix, mgCST 4, 16, 23, 26 and 27 had low scores with mgCST 27 having 40% (8/20) (**Figure 4**). Of the microbiomes with mgCST 27 and scores <0.5, 93.8% (15/16) of vaginal and 100% (6/6) of endocervical were classified as subCST IV-E (**Table S4**), which has a high to moderate abundance of *Prevotella* spp. with *P. bivia* as the most abundant and appear to be unique to Pacific Islanders. This relative abundance profile is not represented in any mgCST.

**Figure 4.**
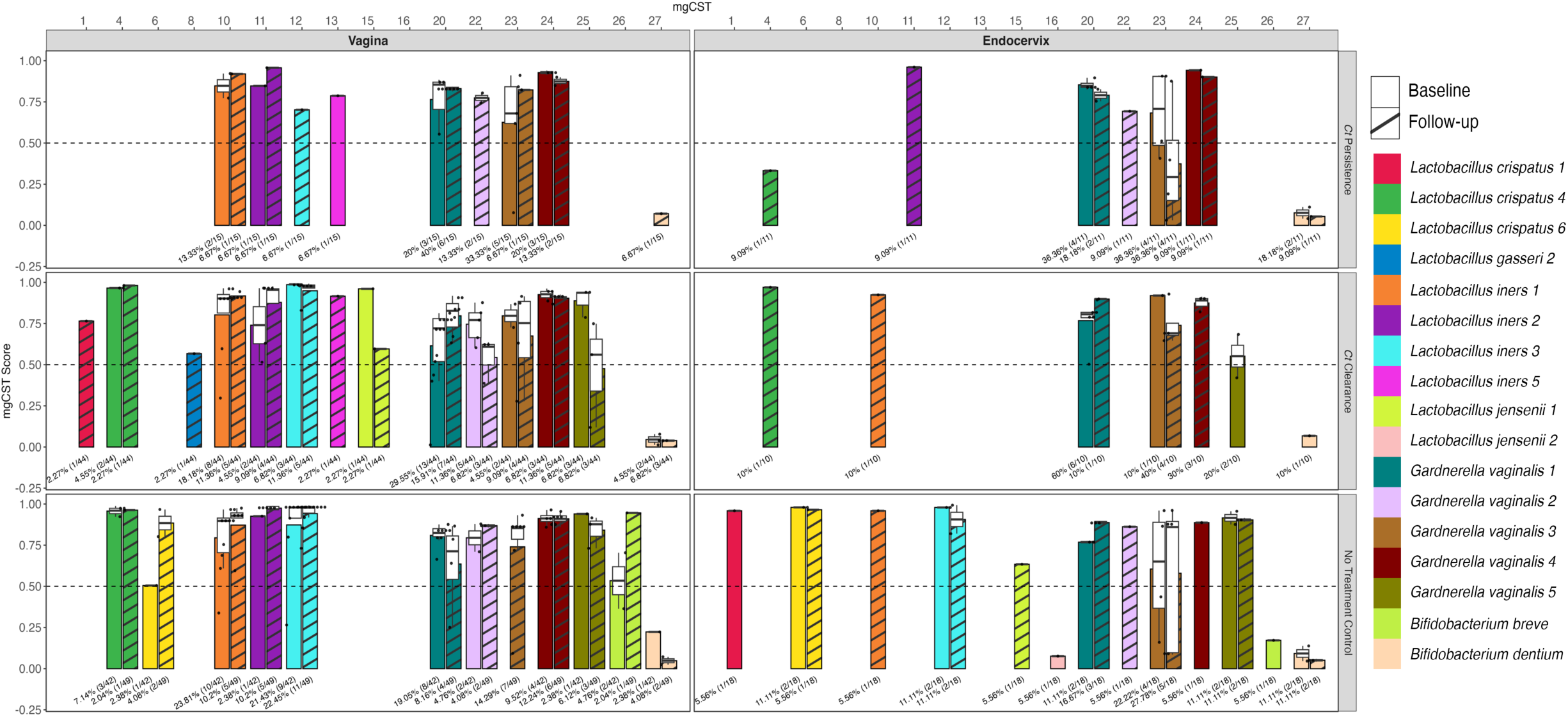
Distribution of mgCST and mgSs types and scores for the paired endocervical (right) and vaginal (left) microbiomes for all three cohort datasets with baseline and follow up shown side-by-side. mgCST classification numbers are shown at the top along the x-axis; mgCST score is shown on the y-axis. 16 (59.3%) of the 27 mgCSTs were found among the Fijian vaginal microbiomes and 15 (55.6%) in the endocervix. The bacterial species are represented by different colors in the color code displayed on the right of the bar graph; no strain types for mgSs are provided by the program, only numbers. The dashed horizontal line across each cohort graph denotes the mgCST score of 0.50. The white box, if shown, within each colored bar of the box plot represents the first (Q1) to the third (Q3) quartile with the median mgCST score denoted by a horizontal line; the whiskers indicate variability outside Q1 and Q3 where the minimum and maximum whisker values are calculated as Q1/Q3 -/+ 1.5 * IQR, with outliers shown beyond the whiskers. The black dots represent individual samples. The numbers below each box plot represent the percentage of samples for that mgCST with the actual numbers provided in parentheses.

There were no statistically significant differences between baseline and follow-up mgCSTs for either vaginal or endocervical microbiomes.

Extending the analysis to metagenomic subspecies (mgSs), in the vaginal microbiomes, *L. iners* mgSs 3 (mgCST 12) was significantly associated with absence of BV (*P*=0.016), while *G. vaginalis* mgSs 4 (mgCST 24) was associated with BV (*P*=0.025) (**Supplementary Table S5**); mgSs 4 was not included in the classification system^31^ as it occurred in relatively low numbers and therefore could not be tested in our samples. *L. iners* mgSs 3 (mgCST 12) and *G. vaginalis* mgSs 1 (mgCST 20) were associated with vaginal *Ct* (*P=*0.042 and *P*=0.026, respectively). *G. vaginalis* mgSs 1 (mgCST 20; *P*=0.007) was associated with *Ct* endocervical infection (**Supplementary Table S5**).

For subCSTs, subCST III-A was associated with absence of BV (*P=*0.006) in the vagina (**Supplementary Table S6**), while subCST IV-A and IV-B were associated with *Ct* (*P*=0.026 and *P*=0.029, respectively). There were no associations of any subCST with *Ct* infection in the endocervix.

### Distinct phylotypes occur in endocervical, vaginal and rectal microbiomes where within host bi-directional genito-rectal and recto-genital species transmissions are evident

Three distinct phylotypes were identified that were similar among the three cohort datasets based on Bray-Curtis hierarchical clustering with each phylotype dominated by a specific group of bacterial species unique to that clade (**Figure 5**). Phylotype I was dominated by *G. vaginalis*, *Lactobacillus* spp., and a group of pathogens including *Prevotella* spp., *Atopobium vaginae*, *Megasphaera genomosp*., *Sneathia amnii* and BVAB1 that are associated with BV consistent with subCST IVs; Phylotype II was dominated by *L. iners*, *L. crispatus*, *G. vaginalis* and the occasional presence of *Prevotella* spp. and *Finegoldia magna*. These two phylotypes were found primarily among endocervical and vaginal microbiomes with CSTs I, III and V. Phylotype III largely consisted of rectal microbiota, dominated by a group of BV-associated and gut bacteria such as *Prevotella* spp., *F. magna*, *Bifidobacterium adolescentis*, *Faecalibacterium prausnitzii* and the occasional presence of *Peptoniphilus lacrimalis*, *Anaerococcus vaginalis* and *Peptoniphilus harei*.

**Figure 5.**
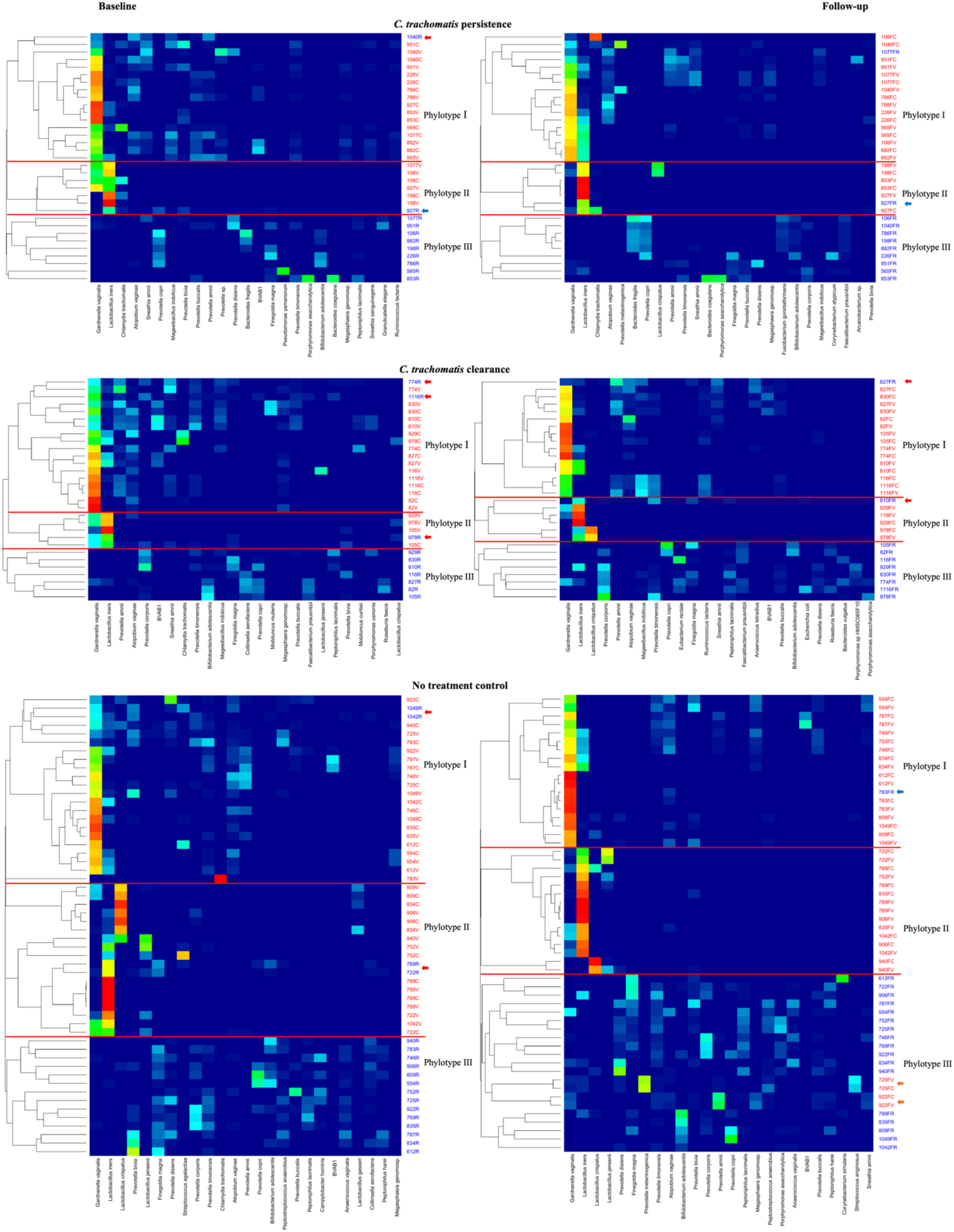
Heatmap based on Bray-Curtis Hierarchical Clustering showing three phylotypes of the paired endocervical, vaginal and rectal microbiomes for each of the three cohort datasets. The level of microbial sharing between microbiomes among the three anatomic sites is shown. The red lines separate the phylotypes within each cohort. The participant ID and the respective anatomic site are shown on the y-axis to the right of the heatmap (i.e., C, endocervix in red; V, vagina in red; R, rectum in blue; F, follow-up). The associated species are shown below each heatmap. Within-host genital to rectal (GRT; rectal samples in Phylotypes I or II; red arrows) or rectal to genital (RGT; endocervical or vaginal samples in Phylotypes III; orange arrows) transmission for each of the three cohort datasets at baseline and/or at follow-up can be appreciated (see also Figure 6). The blue arrows denote complete or almost complete GRT transmissions of *L. iners* or *G. vaginalis*. The dendrograms were set at a distance threshold of 0.9 (see Methods).

The presence of any rectal sample in Phylotypes I and II was indicative of genital to rectal transmission (GRT; **Figures 5, 6**: red and blue arrows) while the presence of any endocervical or vaginal samples in Phylotype III was indicative of rectal to genital transmission (RGT; **Figures 5, 6**: orange arrows). RGT transmission occurred only in two women from the no treatment control cohort in both the vagina and endocervix (**Figures 5, 6**: orange arrows: 922FV, 922FC, 725FV, and 725FC). In some women who were treated, transmissions occurred at baseline and follow-up while for others it occurred at one time point. A complete or near complete GRT resulted in super-colonization of the rectal microbiome in only two women with *G. vaginalis* or *L. iners* (**Figures 5, 6**: blue arrows: 783R and 927R). Some transmissions were partial with the presence of one or two species such as *L. iners* and/or *G. vaginalis* in low abundance in the rectum and were usually present in rectal Phylotype III. Other rectal samples with GRT were present in Phylotypes I and II (**Figures 6**: red arrows). In all cases of complete and partial GRT, the rectal microbiomes could not be classified by the enterotype classification system^32^ because of the mixed origins of the microbial composition of these microbiomes (i.e., enterotypes labelled as “N”; **Figure 6**: red and blue arrows).

**Figure 6.**
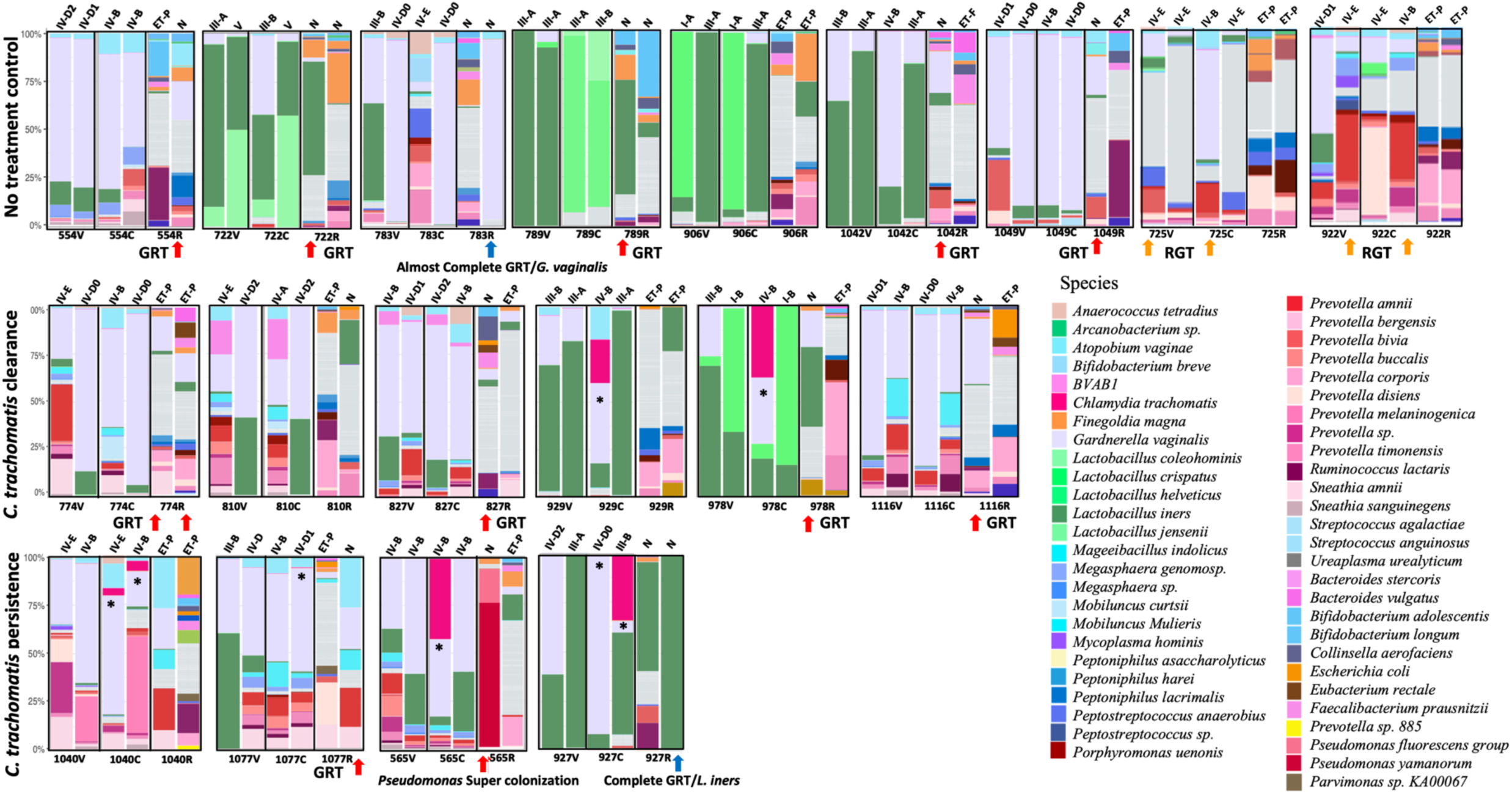
Taxonomic profiles of the paired vaginal, endocervical and rectal microbiomes for each cohort dataset based on the relative abundance at the species level for women exhibiting endocervical/vaginal to rectal transmission (GRT; red and blue arrows) of bacteria and vice versa (RGT; orange arrows). The two bargraphs above the participant ID anatomic site of C (endocervix), V (vagina), or R (rectum) represent baseline (left bargraph) and follow-up (right bargraph) time points. The subCST classification (see **Supplementary Table S4**) is noted at the top of each baseline and follow-up taxonomic bar graph for the vagina and endocervix; enterotype is shown at the top for the rectum where an N denotes an inability to determine an enterotype. The top 25 most abundant species are shown for each cohort dataset (color code shown on right). The remaining species are grouped in the category “Others.”

A relatively high abundance of *Ct*—up to 52%—was visibly noted in baseline endocervical microbiomes of women undergoing antibiotic treatment for *Ct* infection and in the follow up endocervical microbiomes in *Ct* persistence cases (**Figure 6**; see asterisks). The relative abundance profiles of each anatomic site-specific microbiome are shown in **Figures S4-S6**.

### Microbiome function-level sharing between the vagina, endocervix and rectum were different for the Ct persistence, Ct clearance and control cohorts

In comparing the functional determinants across the three anatomic sites, the rectal site was metabolically more diverse with almost twice the number of pathways compared to the endocervix and vagina (**Figure 7**). Moreover, most of the metabolic pathways in the endocervix and vagina were shared with the rectum, either individually or combined.

**Figure 7.**
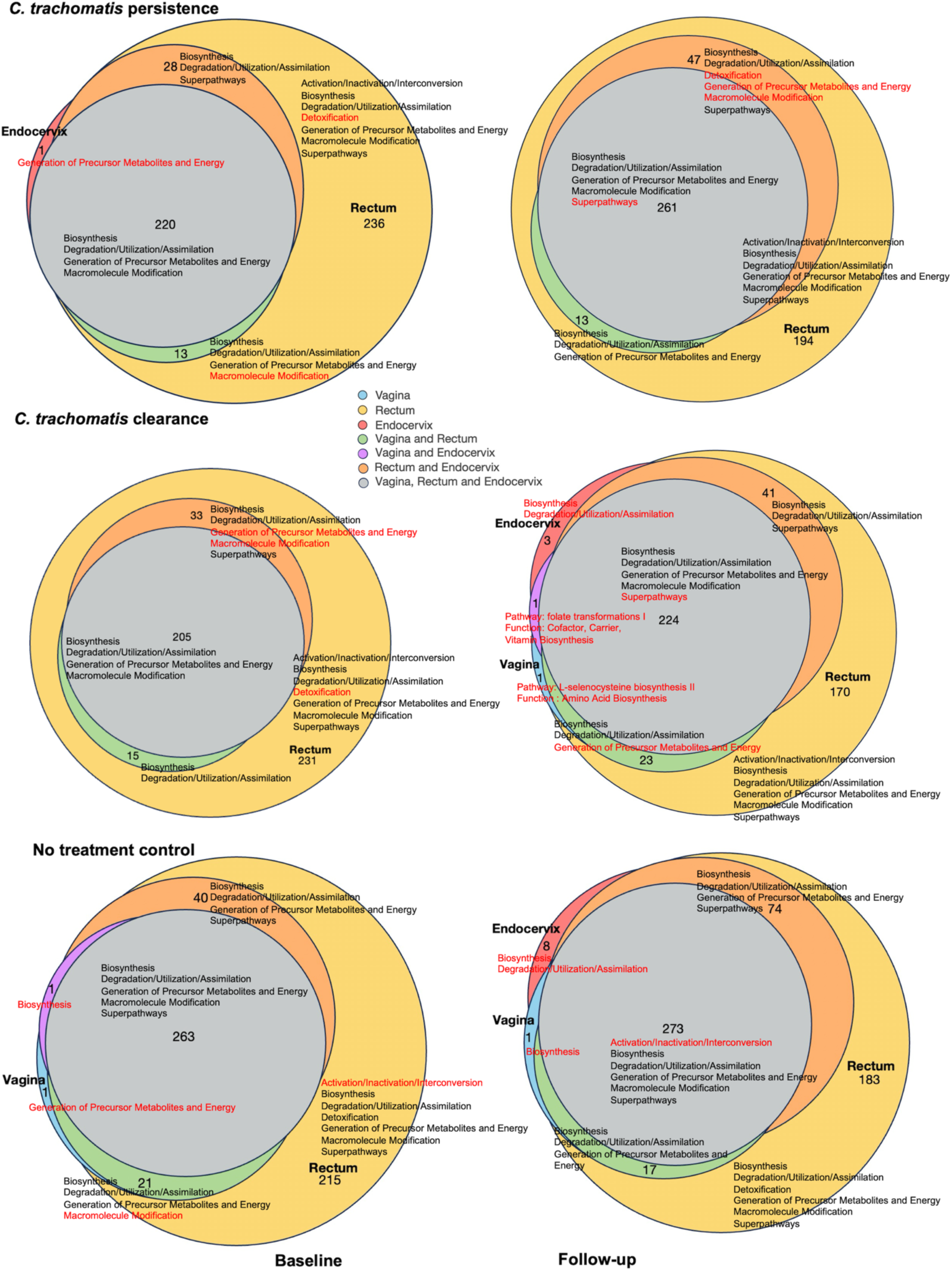
Venn diagram representation of functions and function-level similarities and differences among the paired vaginal, endocervical and rectal microbiomes for the three cohort datasets at baseline and follow-up timepoints. The size of the circle represents the number of function-level pathways comprising that circle. **Supplementary Tables S7 to S12** show the pathways associated with each function for each anatomic site and whether pathways are shared among two or three sites. Vaginal, endocervical and rectal sites are represented as blue, red and yellow, respectively; overlap between two or all three sites are represented in the color key. Red font denotes functions and pathways unique to a particular anatomic site or shared anatomic site.

Among women with *Ct* persistence, only a few pathways were unique to the baseline time point compared to follow up (**Figure 7**; **Supplementary Table S7**) including *Ct*-specific galacticol and phosphatase pathways in the endocervix and rectum. At follow-up, compared to baseline only galactitol degradation in the rectum; glucose and glucose-6-phosphate degradation in the vagina and rectum; and nucleotide degradation in the rectum and endocervix were observed (**Figure 7; Supplementary Table S8**).

Among women with *Ct* clearance, *Ct-*specific pathways at baseline compared to follow up were identical to those for *Ct* persistence with a few that were unique: starch degradation III pathway in the rectum; NAD salvage pathway II in the vagina and rectum; L-lysine and nucleotide biosynthesis in the endocervix and rectum; and stachyose degradation pathways in all three sites (**Supplementary Table S9**). At follow-up, several new pathways were observed: polyamine biosynthesis and amino acid degradation in the endocervix; L-selenocysteine biosynthesis in the vagina; folate transformations in the vagina and endocervix; generation of precursor metabolites and energy in the vagina and rectum (**Figure 7; Supplementary Table S10**).

For the no treatment control cohort, the function-level sharing was similar between baseline and follow up time points with minimal differences (**Figure 7, Supplementary Table S11 and S12**).

### Pathways for generation of precursor metabolites, energy and polyprenyl biosynthesis by Ct-associated microbiomes in the endocervix are associated with Ct infection clearance following antibiotic treatment

Among women who cleared their infection, the function-level community analysis for relative abundance showed a significant increase in carboxylate degradation (*P*<0.05) and decrease in pentose phosphate pathways (*P*<0.05) and polyprenyl biosynthesis (*P*<0.05) at follow-up compared to baseline, but only in the endocervix (**Figure 8A**). **Figure 8B** shows the microbial species associated with the pathways of these functions. Significant pathways at baseline compared to follow up were associated with *Ct* and a few BV-associated bacteria in the endocervix prior to treatment (**Figure 8C**; **Supplementary Table S13**). Similar to the function-level analysis, *Ct*-associated pentose phosphate pathways and polyisoprenoid biosynthesis via MEP pathways were significantly higher at baseline and negligible or absent at follow-up. In addition, isoprene and heme b (aerobic and anaerobic) biosynthesis were also significantly higher at baseline. In the rectum, most pathways were associated with BV-associated and gut bacteria.

**Figure 8.**
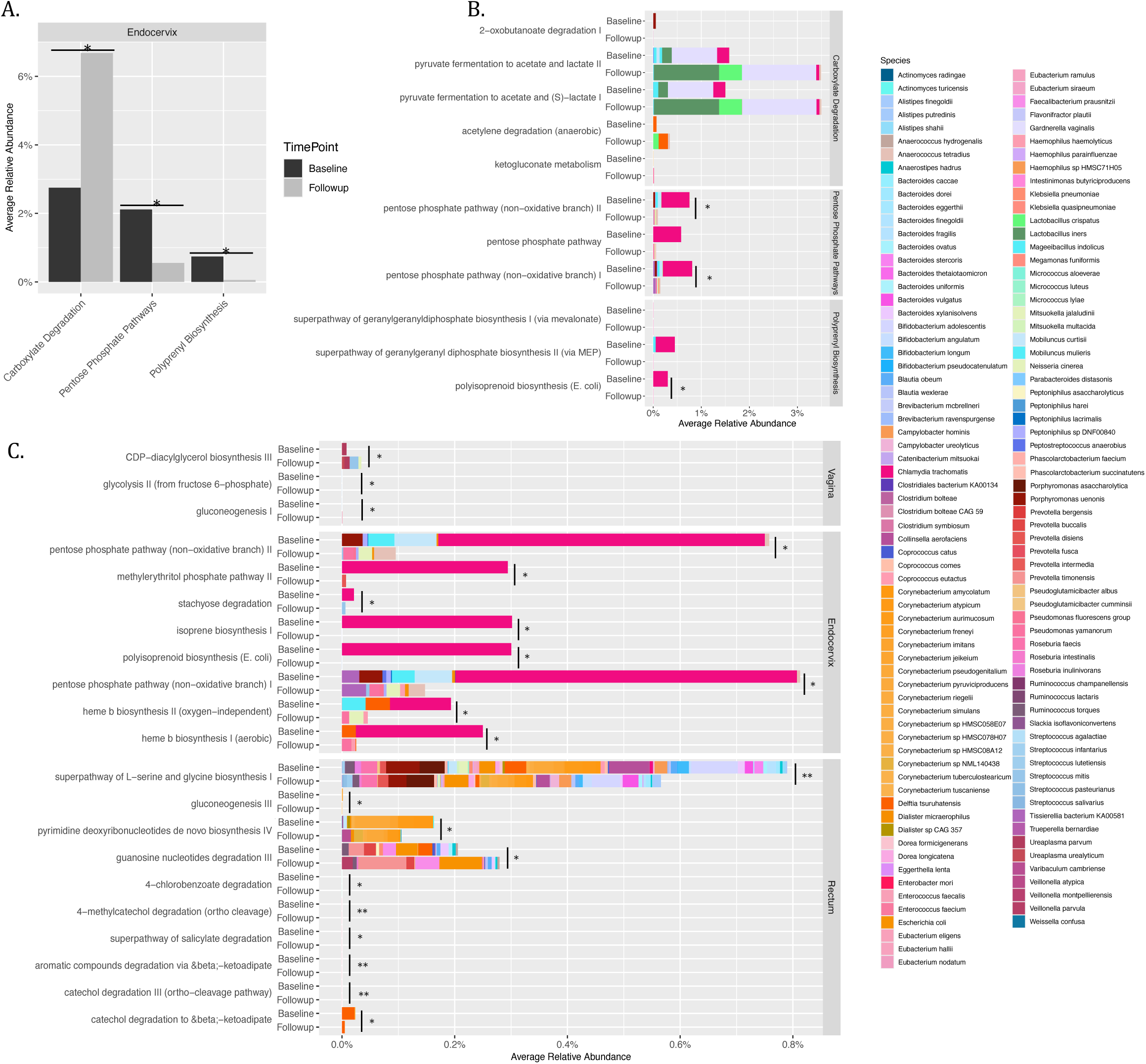
Function- and pathway-level community analysis of paired vaginal, endocervical and rectal microbiomes of women who cleared their *Ct* infection following antibiotic treatment. A. Function-level analysis shows the three functions that had significantly different relative abundances between baseline and follow-up microbiomes for the endocervix. No functions were significantly different for the vaginal or rectal sites. B. Microbial contributions for each function and associated pathways are shown for baseline and follow-up. The species associated with each pathway are in **Supplementary Table S13**. C. Pathway-level community analysis for pathways for each anatomic site that have a significant difference in relative abundance between baseline and follow-up. Species represented in the graphs are color-coded as per the columns to the right of the graphs. *, *P*<0.05; **, *P*<0.005

### Pathways for production of secondary metabolites by Ct, BV-associated and gut bacteria in the rectum act as a hallmark of Ct survival and persistence following azithromycin treatment

Function-level community analysis for the *Ct* persistence cohort revealed a significant increase in secondary metabolite biosynthesis relative abundance following treatment compared to baseline but only in the rectum (**Figure 9A**). **Figure 9B** shows the microbial species associated with the function pathways. Pathway-level community analysis revealed a significant increase in the abundance of pathways related to fatty acid and lipid biosynthesis at follow-up largely driven by BV-associated and gut bacteria; only the CDP-diacylglycerol biosynthesis pathway was associated with *Ct* (**Figure 9**C; **Supplementary Table S14**). Saturated and unsaturated fatty acid, cofactor, and nucleotide biosynthesis pathways were significantly reduced in abundance at follow-up. Among these, 5Z-dodecenoate biosynthesis, palmitoleate biosynthesis I and fatty acid elongation were associated with *Ct* and other BV-associated and gut species.

**Figure 9.**
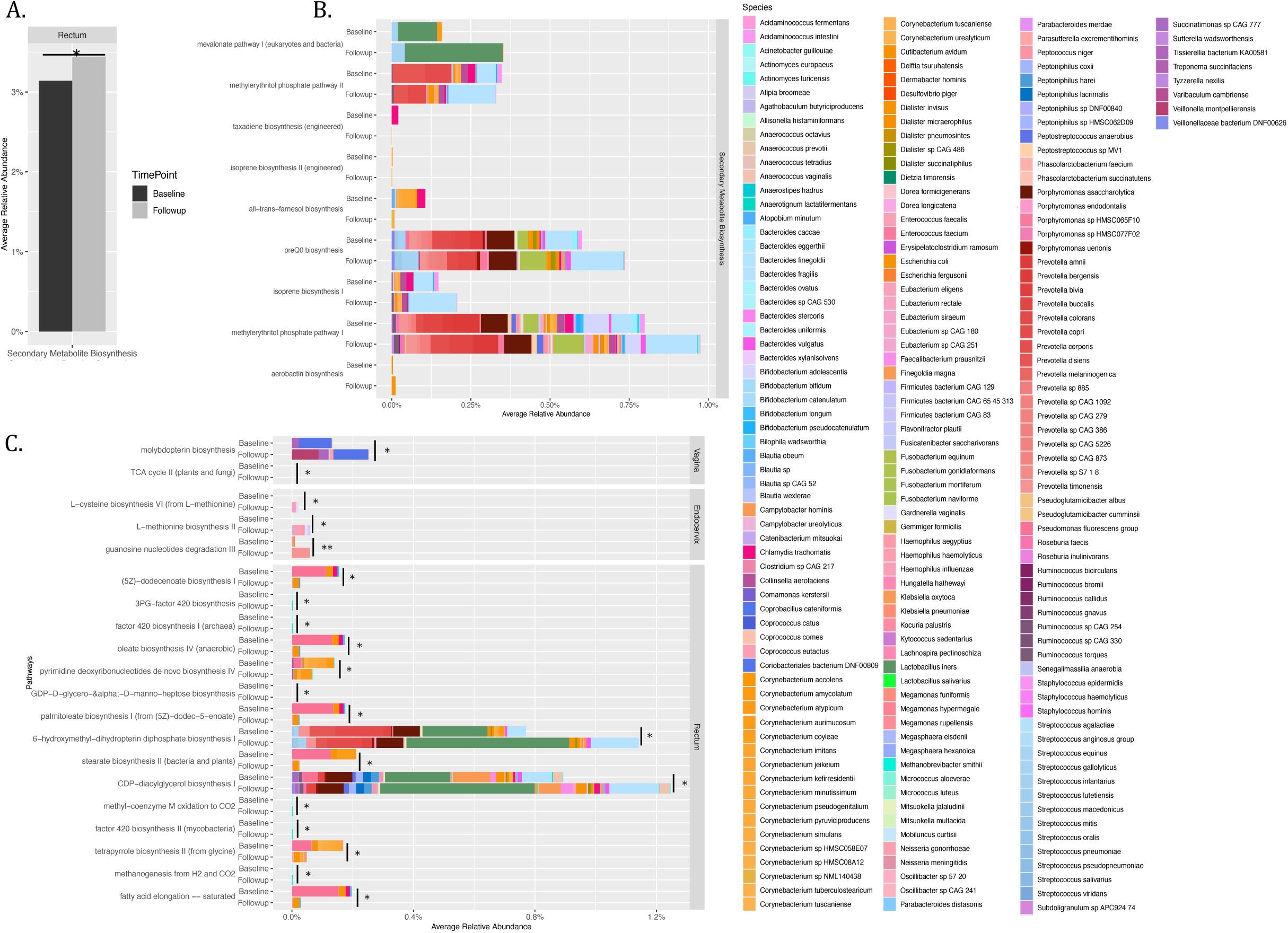
Function- and pathway-level community analysis of vaginal, endocervical and rectal microbiomes of women who did not clear their *Ct* infection following antibiotic treatment. A. Function-level analysis shows one function that had a significantly different relative abundance between the rectal baseline and follow-up samples. No functions were significantly different for the endocervical or vaginal sites. B. The microbial contribution for the function and associated pathways are shown for baseline and follow-up. The species associated with each pathway are listed in **Supplementary Table S14**. C. Pathway-level community analysis shows those pathways for each anatomic site with a significantly different relative abundance between baseline and follow-up. Species represented in the graphs are color-coded as per the columns to the right of the graphs. *, *P*<0.05; **, *P*<0.005

Function-level analysis and associated pathways by microbial species of the no treatment control cohort are shown in **Supplementary Figure S7A** and **S7B**. Only a few pathways were significantly different between the two timepoints but with extremely low pathway abundance (<0.1%) except for a few vaginal, endocervical and rectal pathways driven by gut bacteria (**Supplementary Figure S7C; Supplementary Table S15)**.

### Antibiotic resistance genes (ARGs) and mutations conferring macrolide resistance in Ct, L. iners and G. vaginalis strains from azithromycin treatment and tetracycline resistance in L. iners

We used the CARD (Comprehensive Antibiotic Resistance Database) database to identify any gene with at least a BLAST bitscore of 500 to antibiotic resistance genes (AGRs) among the microbiomes in the three cohort datasets. We also pulled *Ct* 23S rRNA, *rpl*V and *rpl*D genes— known to be involved in azithromycin/macrolide resistance^32–35^—to identify any mutations associated with resistance. AGRs identified by CARD were *erm*B and *erm*X that confer macrolide resistance, and *tet*M that confers tetracycline/doxycycline resistance^36–38^. **Supplementary Table S16** shows the distribution of samples at baseline and follow-up with ARGs and gene mutations conferring the respective resistance for the respective pathogen.

**Figure 10** shows the overall proportion of each ARG and the *rpl*V gene with mutations for the three cohorts. We analyzed the proportion of each gene at baseline compared to follow-up for each of the three cohorts using McNemar’s test (**Figure 10**; see Methods). While there were no mutations in *Ct* 23S rRNA or *rpl*D genes, mutations in the *rpl*V gene were highly prevalent at baseline and follow-up in the vagina and endocervix (74% to 80% for each, respectively), and 25% in the rectum for both timepoints, for the *Ct* persistence cohort (**Figure 10**). These mutations were significantly more likely to occur in vaginal, endocervix and rectal microbiomes (*P*=0.041, *P*=0.013 and *P*=0.007, respectively) in the *Ct* clearance only cohort at baseline compared to follow-up (**Figure 10; Supplementary Table S16)**. Three nonsynonymous mutations in *rpl*V— Gly52->Ser, Arg65->Cys, Val77->Ala—were identified based on MAFTT alignments with a reference gene (see Methods) that have previously been attributed to azithromycin resistance or reduced *in vitro* sensitivity^35^ (**Supplementary Figure S8).** One endocervical sample initially had the Gly52->Ser and Val77->Ala mutations and acquired Arg65->Cys following treatment.

**Figure 10.**
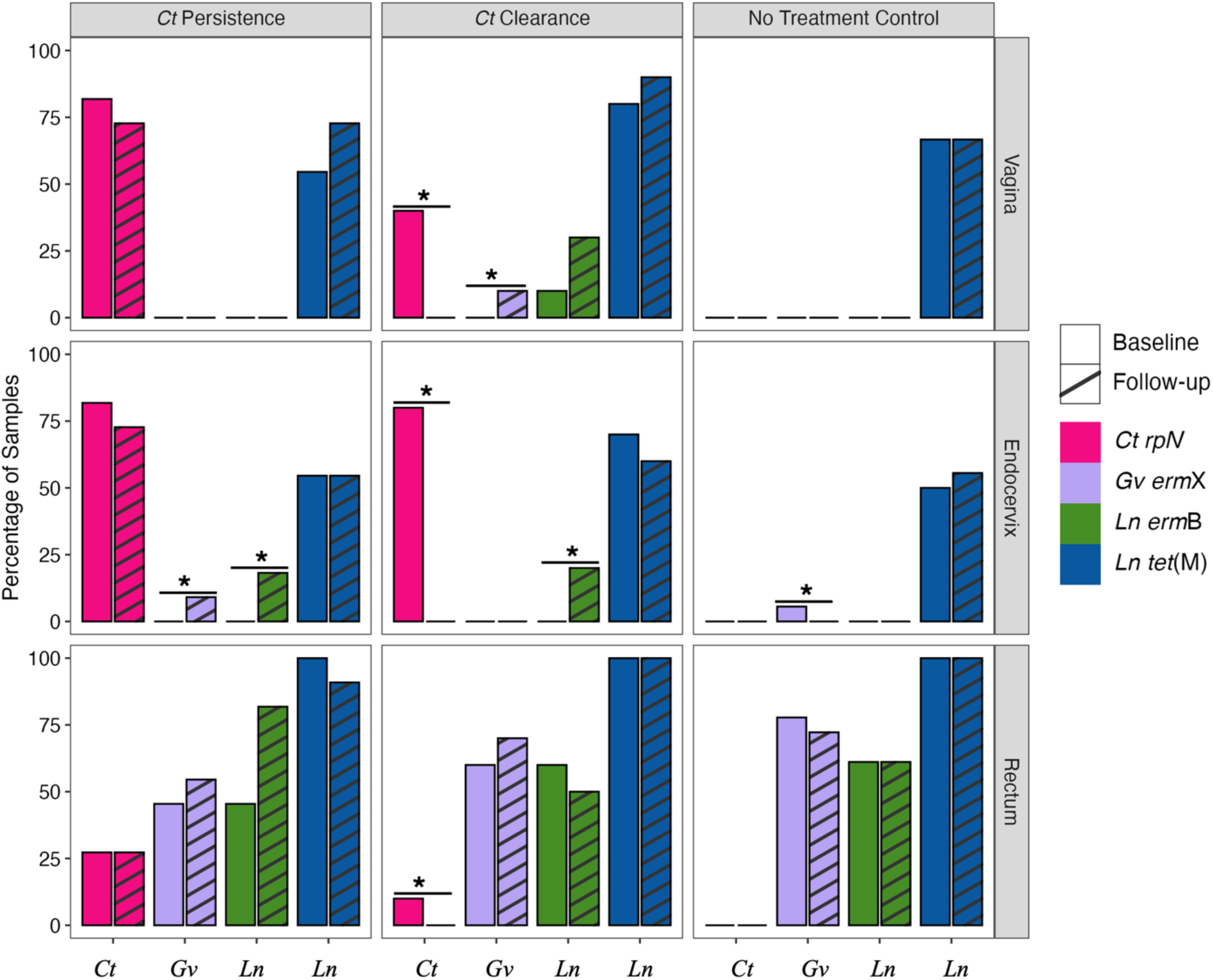
Prevalence of antibiotic resistance genes (ARGs) and mutations associated with *Ct*, *L. iners* (*Ln*) and *G. vaginalis* (*Gv*) based on cohort, anatomic site, and time point. Bar plots show the percentage of samples positive for each gene at baseline and follow-up. Asterisks indicate statistically significant differences between baseline and follow-up within each of the *Ct* persistence, *Ct* clearance, and No treatment cohorts. *, *P*<0.05.

The presence of the *erm*B gene in *L. iners* was significantly associated with endocervical microbiomes in the *Ct* clearance (*P*=0.043) and *Ct* persistence (*P*=0.027) cohorts following azithromycin treatment compared to baseline (**Figure 10**). The presence of *erm*B genes was confirmed by pulling the designated CARD reads and aligning the respective *L. iners erm*B genes from the two cohorts with a reference *erm*B gene (**Supplementary Figure S9**). There was also a high prevalence of the *tet*M gene, which confers tetracycline resistance, in *L. iners* associated with all anatomic sites for the three cohorts (**Figure 10**). *L. gasseri* and *L. crispatus* were also evaluated but no macrolide-specific ARGs were identified.

The *erm*X resistance gene was found in *G. vaginalis* among all cohorts, but primarily in rectal microbiomes at baseline and follow-up (**Figure 10**), confirmed by sequence alignments with the reference gene (**Supplementary Figures S10**). There was a significant increase in this gene in the endocervix of the *Ct* persistence cohort (*P*=0.009) at follow-up, the vagina in the *Ct* clearance cohort (*P*=0.016) at follow-up, and the endocervix in the control cohort (*P*=0.000) at baseline (**Figure 10**).

## Discussion

We previously evaluated microbial diversity and community composition among paired endocervical and vaginal microbiomes of Pacific Islanders in Fiji and discovered that over 35% were unique and could not be classified using the subCST system developed in 2020^17^. Many microbiomes were diverse with a high relative abundance of facultative and strict anaerobic organisms such as *G. vaginalis* and *Prevotella spp*. that comprise subCSTs III and IV. We therefore developed subCSTs IV-D0, D1, D2 and E to capture the diversity and microbial composition of these microbiomes^16^. Here, we investigated whether the new mgCST and mgSs classification system^31^ could better define the microbiomes and reveal antibiotic effect modifications. Over 10% of vaginal and >25% of endocervical microbiomes had low similarity scores (i.e., <0.5) to the closest mgCST centroid that frequently was mgCST 27, which comprises a catch-all of bacteria that are not defined. Samples in this category tended to match well with subCSTs IV-D0, IV-E or other IVs. Further, while we found some significant associations of, for example, *L. iners* mgSs 3 with absence of BV, which was the same for subCST IIIA, mgSs are not defined at the strain level and therefore can’t be used to further explore their association with metadata or other microbiome data. Consequently, we used the expanded subCST classification system that we developed^16^.

Comparing the effect of treatment on CSTs, only one woman in the *Ct* clearance cohort transitioned to a ‘healthy’ subCST I-B in vaginal and endocervical microbiomes. Among women with persistent *Ct*, only two women shifted from subCST IVs to subCST IIIs, dominated by *L. iners.* While Pacific Islander microbiomes tend towards more diverse and BV-associated species^16,28^, azithromycin treatment showed no improvement in subCST or BV, similar to recent 16S rRNA studies of vaginal samples^20,22^.

The lack of improvement in the microbiomes following treatment indicates that the risk of perpetuating *Ct* infection and acquiring other STIs remains. We found that women in the *Ct* persistence and clearance cohorts were significantly more likely to have hrHPV types in the endocervix post-treatment compared to uninfected, untreated women. Many hrHPV types were the same as those at baseline. Previous studies have shown that *Ct* increases susceptibility to HPV infection and interferes with its clearance, thereby promoting persistence^39,40^. Potential mechanisms include epithelial cell damage that facilitates invasion and depression of the host immune response^40,41^. In the latter case, *Ct* decreases effective antigen presentation by dendritic cells, alters the density and antigen-presenting ability of Langerhans cells, and induces T lymphocyte apoptosis with loss of CD4+ and CD8+ T cells that limits cell-mediated immunity, favoring HPV persistence^40,42^. These and other studies have also indicated that co-infection with *Ct* can potentiate progression to cervical cancer ^43,44^.

In addition to hrHPV, *Ng* increased in all sites at follow-up for the *Ct* persistence cohort but was significantly higher only in the endocervix compared to controls. Using LDM, we found that, in addition to *Ng,* other microbiota were significantly associated with *Ct* persistence in the rectum. These included the aerobic and anaerobic bacteria *S. lutetiensis*, *Prevotella* spp., *Ruminococcus* sp., and *F. motiferum*. Both *S. lutetiensis* and *Fusobacterium* spp. are associated with ulcerative colitis (UC) and colorectal cancer ^45–49^. The latter species have increased virulence when other aerobic and anaerobic pathogenic organisms of the oral and gastrointestinal tract are present^47,50–52^. Collectively, these organisms may disrupt the rectal mucosa and promote an environment that predisposes to persistent inflammation and infection by STIs such as *Ct, Ng* and HPV where tissue penetration of antibiotics is limited. Additional studies in this area are warranted to better understand pathogenic mechanisms that can inform appropriate treatment modalities.

We also evaluated whether specific bacteria at baseline could predict *Ct* persistence at any anatomic site at follow-up. While none were found for the endocervix or vagina, the gut-associated bacteria *C. hominis, C. catus, R. callidus* and *R. inulinivorans,* along with a decrease in *L. acidophilus* and *L. fermentum,* served as significant biomarkers of persistence in the rectum. *Campylobacter* spp. have been linked to Inflammatory Bowel Disease (IBD), Crohn’s Disease (CD) or other gastrointestinal diseases^53–55^. *C. catus* is known for playing a role in health via production of vitamin B and acetate, which are inversely correlated with visceral fat, but significantly decreased in UC and IBD compared to controls ^56,57^ as is *R. callidus* in CD and *Roseburia* spp. in CD, UC, and CRC^58^. A decrease in the two *Lactobacillus* spp. could compromise gut health by disrupting the balance between beneficial and harmful bacteria, decreasing digestion and diminishing the host immune response^59^. Their decrease could also limit repopulation of the vagina where they could assist in homeostasis and perhaps modulate BV and its recurrence^60^. These bacteria, or lack thereof, would benefit *Ct* and other STI pathogens and their persistence. Overall, the data suggest the need for a larger prospective study to further investigate potential species that may be predictive of *Ct* persistence.

In our previous pilot study^28^, we found that high *Ct* abundance led to genito-rectal transmissions. In the current study, these transmissions were common regardless of *Ct* infection in all cohorts at both baseline and follow up whereas rectal to genital transmissions occurred only in the control cohort, perhaps due to our small sample size. The former transmissions mainly involved *L. iners* and *G. vaginalis*. Translocation of species from the vagina to the rectum and vice versa are probably more frequent than previously recognized. This is supported by a recent review of inter-anatomic site cross-talk^61^ and a study that used 16S rRNA typing and found similar BV-associated bacteria in both the vagina and rectum of pregnant women in Japan^62^. These transmissions could contribute to the high rates of BV and diverse anaerobic bacteria in our population where the rectum acts as a reservoir that enhances non-*Lactobacillus* dominated vaginal and endocervical microbiomes. These microbial compositions would maintain or increase the risk of acquiring *Ct* and other STIs in the endocervix and rectum, favoring transmission and re-infection, which may in part explain the current high rates of *Ct* and *Ng* in Fiji ^4,8,27^.

The endocervix, but not the rectum or vagina, had significantly more metabolic pathways at baseline compared to follow up for the *Ct* clearance cohort. *Ct* was primarily responsible for this activity, which included biosynthesis of isoprenoids (via MEP, isoprene and polyisoprenoid biosynthesis pathways), energy production, and heme b metabolism. Metabolites in the MEP pathway are thought to disrupt histone-DNA interactions that then allow for the transformation of chlamydial infectious elementary bodies (EBs) to metabolically active reticulate bodies (RBs) in preparation for replication^63,64^. The production of ATP and reducing power, and metabolism of heme are important for transition from RBs to EBs that are released from the cell to spread within tissue and be transmitted ^65,66^. After treatment, there was a significant drop or absence in isoprenoid biosynthesis for *Ct*, which is not surprising as azithromycin inhibits protein synthesis that occurs at the RB stage ^67^, indicating successful treatment.

In the *Ct* persistence cohort, only the rectum had a significant increase in metabolic pathways between baseline and follow-up and that was for secondary metabolite production related to isoprenoid biosynthesis. *Ct*, BV-associated and gut bacteria contributed to this increase. A few studies provide evidence that isoprenoids or derivatives of isoprenoids allow *Ct* to adapt to the intracellular host environment through enhanced oxidative stress resistance^64,68^. The fact that there were no additional significant differences in metabolism between the two time points suggest that major shifts in metabolism did not occur in response to treatment, which is consistent with persistent infection in the endocervix and rectum. However, it is not clear why some individuals cleared their infections while others did not. It is possible that, in the persistence cohort, the rectum is the primary site of infection, which is known to be difficult to treat—only ∼70% of rectal infections are effectively treated with azithromycin^69–71^. Microbial exchange with the vagina/endocervix may therefore be more frequent in this cohort. At baseline, *Ct*, *Prevotella spp*. and *Corynebacterium spp*. contributed to fatty acid (FA) biosynthesis pathways. Following treatment, there was a significant increase in phospholipid biosynthesis by *Ct, L. iners* and other gut-related bacteria including *Prevotella spp*., *Porphyromonas spp*., *Bacteroides spp*. Experimental evidence suggests that *Ct* encodes all the genes to endogenously synthesize FAs and phospholipids necessary to assemble its intracellular membrane and for proliferation^72^. Phospholipids also play a crucial role in protection against antimicrobials, which may explain the increase in their synthesis at follow up during *Ct* persistence. Indeed, a recent study found that metabolic reprogramming can occur in *Ct* persistence—induced in this case by β-lactam treatment—causing a switch from the tricarboxylic acid (TCA) cycle to FA biosynthesis^73^.

There are some anecdotal reports of presumed tetracycline, macrolide or multi-drug resistance in clinical *Ct* isolates^74–78^. We explored the possibility that azithromycin resistance may play a role in *Ct* persistence in Fiji. While azithromycin resistance is typically attributed to heterotypic resistance (i.e., a subpopulation of bacteria with lower susceptibility to the antibiotic) or bacteriostatic antibiotic treatment^79^, we found evidence for possible homotypic (genetically inherited) resistance in *Ct rpl*V with three nonsynonymous mutations in the ribosomal L22 protein. These mutations had a high prevalence of ∼80% in the vaginal and endocervical sites with ∼25% in the rectum in the *Ct* persistence cohort with a lower prevalence in the *Ct* clearance cohort, which may in part explain clearance. The macrolide binding site in L22 is immediately downstream of a variable region where the three mutations were located^80^. A study in Japan found the same mutations in three of five male urethritis patients who had persistent infection following treatment with extended-release azithromycin^81^. The minimum inhibitory concentrations (MICs) for azithromycin were all 0.08 μg/ml, which does not explain *Ct* persistence in these men. However, in a study in Russia, three of four patients had these mutations, and two of the three had MICs of >5.12 μg/ml pre- and post-treatment while one patient remained sensitive^33^. The former two had repeated episodes of cervicitis and urethritis, respectively. While these collective findings are suggestive of resistance, larger studies are needed to more fully understand the distribution of potential azithromycin resistance based on these mutations and the mechanism(s) by which resistance may occur.

We also found significant enrichment of *erm*X in *G. vaginalis* in the endocervix of women with *Ct* persistence post-treatment. The *G. vaginalis* strains with *erm*X—JNFY4, JNFY1 and JNFY9—belong to the virulent subtype C clade, which is associated with symptomatic BV and moderate to high biofilm formation^37^. Our previous microbiome network analysis found an association with endocervical *Ct* and *G. vaginalis* interactions supporting *Ct* persistence^16^. Additionally, there was a significant enrichment of *erm*B in *L. iners* following treatment in the endocervix of both the *Ct* clearance and persistence cohorts. A high proportion of *tet*M was identified in *L. iners* in all cohorts in all anatomic sites. This gene is frequently transferred via transposons and is common among gram-negative and gram-positive pathogens, causing tetracycline resistance by encoding a protein that prevents binding of tetracycline to ribosomes^38^. The emergence of resistance to the two main antibiotics used to treat *Ct* following treatment coupled with the increase in relative abundance of *L. iners* and *G. vaginalis*, which are risk factors for *Ct* infection and reinfection in addition to other STIs^19,82,25,26^, are a major concern. This highlights the potential collateral effect of antibiotics on microbial ecosystems where there is an increase in abundance of non-target pathogens that increase the risk of re-infection or persistence of the target pathogen, in this case *Ct*, in addition to other STIs.

Our results reveal perturbing effects of azithromycin treatment with increased risk of hrHPV and *Ng* co-infections, macrolide and tetracycline resistance among *L*. *iners* strains following treatment that can also promote their growth and increase the rick for other STIs, and mutations in *Ct* that may confer azithromycin resistance, indicating the need for further prospective studies that may elucidate the depth and breadth of resistance to azithromycin, and also reveal novel therapeutic strategies to treat *Ct* and protect and restore the microbiome.

## Methods

### Study design and patient characteristics

Women aged 18 to 40 years who were seen at Fiji MHMS Health Centers and two university clinics in the Central Division, Viti Levu, Fiji, were consecutively enrolled after informed consent as a convenience sample and followed quarterly for 12 months as part of the parent study^4,16^. The research was approved by the Institutional Review Boards of the MHMS and the University of California San Francisco.

Study participants and their clinical, microbiologic, and sample data were anonymized using unique ID numbers. Data supplied to this study for participants seen at baseline and follow up visits included age, ethnicity/race, signs and symptoms, presence or absence of BV based on Amsel Criteria^43^ and results of the Cepheid Xpert® CT/NG diagnostic assay (Cepheid, Sunnyvale, CA). Women who tested positive for *Ct* at their baseline visit were treated with a single oral dose of 1g Azithromycin. Paired endocervical, vaginal and rectal samples were available for 26 women who were infected with *Ct* at baseline and either cleared their infection (n=15) or remained infected (n=11) at follow up and 18 women who remained uninfected at both time points (**Figure 1**).

### Sample processing, metagenomic shotgun sequencing, pre-processing and taxonomic analysis

About 400µl of each clinical sample was treated with a lysis cocktail containing 100µl lysozyme, 12µl mutanolysin, and 6µl lysostaphin as we described^28^. DNA was purified using the QIAamp DNA Mini Kit (Qiagen, Germantown, MD) and quantified using the Qubit^TM^ HS assay kit (Invitrogen, Carlsbad, CA) according to manufacturer’s instructions.

Genomic DNA (120 ng per sample) was prepared using Illumina Nextera XT kits and subjected to metagenome shotgun sequencing using 150bp paired end reads on an Illumina Novaseq platform including a Zymo artificial controls with all reagents except for DNA. Sample preprocessing was performed to remove adaptors and human DNA as previously described^28^. Data from vaginal and endocervical samples were taxonomically profiled using VIRGO^83^. Rectal samples were profiled using Metaphlan v3.0.14^84^. The presence of *Ng*, *Mg*, *Tv* and *Candida* were identified and quantified from the VIRGO endocervical and vaginal data as previously described^28^. High risk (hr) and low risk (lr) Human Papilloma Virus (HPV) types were identified using HPViewer^85^ for all three anatomic sites.

### subCST and mgCST classifications for endocervical and vaginal microbiomes, and enterotype determination for rectal microbiomes

Endocervical and vaginal subCSTs were identified using the nearest centroid classifier algorithm in VALENCIA^17^ in addition to our in-house subCST classification system that includes the Pacific Islander microbiomes as described^16^. We also classified our metagenomes using the new mgCSTs and mgSs system^31^. mgSs refer to the combination of strains of a specific species present in an mgCST that may be associated with, for example, BV. mgCSTs include assignment of a similarity score to a mgCST centroid of which there are 29. A score of <0.5 indicates low similarity to the assigned centroid.

In addition to the three cohort databases described above (**Figure 1**), an additional larger vaginal cohort dataset—comprised of 34 women who were *Ct*-positive and either remained positive (n=5) or became negative (n=29) after treatment along with 24 women who were *Ct*-negative at both time points—was also used to determine subCST, mgCST and mgSs classifications.

Rectal microbiomes were classified based on the enterotype method developed for gastrointestinal microbiomes as described^28,86^.

### Microbial Functional Determinants and Resistome analysis

Functional modules of the endocervical, vaginal and rectal microbiomes at the community- and species-levels were identified using HUManN2 v3.0^87^. Additionally, protein and pathway attributes generated through VIRGO for vaginal and endocervical microbiomes were also analyzed. Any unintegrated and unmapped data were excluded from analysis. The resulting reads were then transformed into the relative abundance of the reads per sample and subsequently visualized (see below).

Megahit^88,89^ was used to generate contigs from the MSS files, which were run through the resistance gene identifier (RGI)—an ARG detection model available in CARD^90^. Only perfect—an ARG with all amino acids matching the CARD database—and strict hits defined as a hit to an ARG with at least a bit score of 500, were used; loose hits were discarded. AGRs specific to *L. iners*, *L. crispatus*, *L. gasseri* and *G. vaginalis* were extracted from CARD outputs in addition to 23S rRNA, *rpl*D and *rpl*V genes from *Ct*. To further confirm the presence of ARGs and any associated mutations, contigs specific to macrolide resistance genes were extracted from the samples and the sequences were aligned using MAFFT plugin in Geneious Prime 2024.0.3 (**Supplementary Figures S8-S10**). CARD models of the associated genes and their species were used to extract information related to the literature, strains harboring resistance genes and the respective protein homolog model, which were then integrated into the alignments and data interpretation.

### Statistical Analysis and Visualization

Proportional comparisons of the hrHPV types across the three anatomical sites were analyzed using Fisher’s Exact or Chi-squared Test. These tests were applied at baseline and follow-up timepoints within each of the three cohorts. The likelihood of maintaining the same hrHPV at baseline and follow-up, within each cohort, was tested using McNemar’s Chi-squared Test. The One-Sample Proportion Test was also performed to compare the proportion of samples with hrHPV at baseline and follow-up across all three cohorts. A significance level of *P*=0.05 was used for each analysis (**Supplementary Table S3**).

The mgCST and mgSs designations and associated scores were determined for the paired endocervical and vaginal cohort datasets (**Figure 1**) and the larger vaginal cohort dataset using the relative abundance of each sample as described^16^ (**Figure 4**). The mean mgCST score for each mgCST was calculated, and the white boxplots were overlaid on the bar plot to show the distribution of the data as first quartile (Q1), median, and third quartile (Q3) for each mgCST. The whiskers represent the minimum and maximum values, calculated as Q1/Q3 ± 1.5 times the interquartile range (IQR). Any values outside of this range are displayed as outliers. The ggplot package in R ^91^ was used to generate the figure.

To examine the association of BV or *Ct* with mgCST (**Supplementary Table S5**) and with subCST (**Supplementary Table S6**) designations, a chi-square test was used to compare the number of positive and negative women across each classification system. For example, the expected number of *Ct* positive women was calculated using the formula (total *Ct*+ * no. *Ct*+)/total no. women, and similarly for *Ct* negative women. The CHISQ.TEST function in excel was used to calculate the chi-square value using the observed and expected values.

Microbiome analysis was performed using R software packages ^91^: Vegan (v.1.4-5)^92^, Adespatial (v.0.3-14)^93^, ggplot2 (v.3.3.5)^91^, CGPfunctions (v.0.6.3)^94^, FSA (v.0.9.3)^95^, ggpubr (v.0.4.0)^96^, dendextend (v1.14.0)^97^, dplyr (v.1.1.4), and stats (v.4.4.0) to create relative abundance plots (**Figure 6; Supplementary Figures 4 to 6**). The transition from one subCST to another within each dataset from baseline to follow-up were visualized using the CGPfunctions^94^ library (**Supplementary Figure 3**).

A series of analyses were performed on the microbiomes, comparing independent samples between two groups at one time point, such as the *Ct* clearance vs. *Ct* persistence cohorts at baseline or follow-up; comparing paired samples between the two time points, baseline vs. follow-up, within a cohort, such as the baseline to follow-up in *Ct* clearance cohort; and comparing baseline-to-follow-up changes between anatomic sites or two cohorts (e.g., women who resolve their infection vs women who remain infected), which are addressed by analyzing the time-site or time-cohort interaction. For each analysis, the microbiome profiles were analyzed in terms of alpha-diversity, beta-diversity, and relative abundances at individual species. The alpha-diversity was measured by the richness and Shannon Index of the species. Each alpha index was compared between groups of independent samples by the Wilcoxon rank-sum test or between paired samples by the Wilcoxon signed-rank test (**Supplementary Figure 1**). The beta-diversity was measured by the Bray-Curtis or Jaccard dissimilarity metric. The resulting distance matrices were used to visualize sample clustering by the principal coordinates analysis (PCoA) plot and tested for shifted clusters along a given covariate (e.g., group label) by the PERMANOVA model^29,98^ for both independent and paired samples (**Supplementary Figure 2**). The relative abundances at individual species were analyzed by the Linear Decomposition Model (LDM)^29^, which can compare relative abundances between groups of individual samples and paired samples, as well as testing time-site or time-cohort interactions (**Figure 3**). The LDM detects significant species while controlling for the false-discovery rate (FDR) at a pre-specified level (20% here). The LDM also provides a global p-value for testing a hypothesis at the community level.

Genito-rectal transmissions and sharing of microbial species between the anatomic sites of each woman within a cohort were analyzed and visualized by Bray-Curtis hierarchical clustering between samples in each cohort and timepoint using Vegan^92^. Dendograms were generated using dendextend^97^ (with a distance threshold of 0.9) and heatmaps were created using ggplot^91^ (**Figure 5**).

For functional attributes, the pathway abundance file from the HUManN2 v3.0^87^ output was used. Microbiota across each function and pathway within the microbial community of the respective microbiomes were compared between baseline and follow-up timepoints across each study cohort to first create a Venn representation using ggVennDiagram^99^—depicts the function-level sharing and uniqueness for each cohort across the three anatomic sites and two time points (**Figure 7**). The paired two-sample T test and paired Wilcoxon test with an alpha significance level of 0.05 was used for the function- and pathway-level analyses. The data were visualized in ggplot^91^ (**Figures 8, 9**; **Supplementary Figure 7**). *Ct*-specific pathway and function-level data for baseline and follow-up time points for the three cohorts are provided in **Supplementary Tables S7 to S12.** The statistically significant differences in functions and pathways between baseline and follow-up in the *Ct* clearance and *Ct* persistence cohort for *Ct* and other bacteria are provided in **Supplementary Tables S13, S14.**

For the resistome analysis, the presence/absence of genes was compared between baseline and follow-up timepoints for each cohort and the anatomic site using McNemar’s test in R package stats (v.4.4.0). Only significant results at *P*<0.05 values were plotted using the R packages^91^ dplyr, tidyr, ggplot2, ggtext, and ggh4x (**Figure 10B**). The *Ct* genes 23S rRNA, *rpl*D and *rpl*V were aligned against reference strain D/UW-3/CX to discern any gene and nonsynonymous amino acid mutations (**Supplementary Figures S8)**. We also compared the data across all three cohorts, their timepoints and anatomic sites to identify genes unique to each cohort. Only genes which appeared in at least 15 microbiome samples were considered and plotted as above. To generate the plot, R packages dplyr (v.1.1.4), tidyr (v.1.3.1), ggplot2, ggtext (v.0.1.2), ggh4x (v.0.2.8) were used.

## Data availability

All scripts generated for this research relating to data visualization and statistical analyses can be found at https://github.com/FijiProspectiveStudy. The MSS FASTQ files for all the rectal metagenomes were submitted to the NCBI-SRA under the BioProject accession number PRJNA1153641 and the endocervical and vaginal metagenomes were submitted to the NCBI-SRA from our previous study^15^ under the accession ID PRJNA982400.

## Acknowledgements

We thank the parent study for providing the deidentified data for this study and Fijian colleagues: Dr. Rachel Devi; Dr. Kinisimere Nadredre; Dr. Mere Kurulo; Dr. Jenni Singh, and Dr. Darshika Balak. This research was supported by Public Health Service grant from the National Institutes of Health R01 AI151075 (to DD and TDR).

## Author contributions

SB: Data curation, Formal analysis, Methodology, Visualization, Writing – original draft, Writing – review and editing | OO: Formal analysis, Methodology, Software, Visualization, Writing – review and editing | YH: Formal analysis, Methodology, Software, Writing – review and editing | RW: Formal analysis, Visualization, Writing – review and editing | MK: Investigation, Supervision, Writing – review and editing | MD: Formal analysis, Visualization, Writing – review and editing | RK: Formal analysis, Visualization, Writing – review and editing | TDR: Conceptualization, Data curation, Funding acquisition, Investigation, Methodology, Resources, Supervision, Writing – review and editing | DD: Conceptualization, Formal analysis, Funding acquisition, Investigation, Methodology, Project administration, Resources, Supervision, Validation, Writing – original draft, Writing – review and editing. SB and OO are “co-first authors”.

