## Supplemental Figures for "Effect of Azithromycin treatment on the microbial composition, functional dynamics and resistomes of endocervical, vaginal and rectal microbiomes of women in Fiji with *Chlamydia trachomatis* infection"

### Supplementary Figures

**Supplementary Figure 1**. Alpha diversity measured by Shannon index and log richness of species for the paired vaginal, endocervical and rectal microbiomes for women without treatment, those who cleared their *C. trachomatis* infection post-treatment, and those who had persistent infection after treatment. *P* values are denoted within each box in the lower righthand corner.

**Supplementary Figure 2**. Beta diversity measured by Jaccard and Bray-Curtis metric for the paired vaginal, endocervical and rectal microbiomes for women without treatment, those who cleared their *C. trachomatis* infection post-treatment, and those who had persistent infection after treatment. *P* values are denoted within each box in the lower righthand corner.

**Supplementary Figure 3**. Chord diagram representation of the transition in sub-Community State Types (subCSTs) for the three cohort datasets for endocervical, vaginal and rectal sites. *Transition from a *Lactobacillus*-dominated microbiome (subCST I, II or V) to a microbiome dominated by *L. iners* or diverse anaerobic and facultative bacteria (subCST III or IV) or vice versa (‡).

**Supplementary Figure 4**. Taxonomic profiles of the vaginal microbiomes, paired with the endocervical and rectal microbiomes (not shown), based on relative abundance at the species at baseline and follow-up for the three study cohort datasets. The participant ID# is shown at the top of the respective box that contains baseline (left) and follow-up (right) bar graphs and the in sub-Community State Types (subCSTs) noted below the bar graphs. The samples exhibiting genito-rectal transmissions are shown in **Figure 6**.

**Supplementary Figure 5**. Taxonomic profiles of the endocervical microbiomes, paired with the vaginal and rectal microbiomes (not shown), based on relative abundance at the species at baseline and follow-up for the three study cohort datasets. The participant ID# is shown at the top of the respective box that contains baseline (left) and follow-up (right) bar graphs and the in sub-Community State Types (subCSTs) noted below the bar graphs. The samples exhibiting genito-rectal transmissions are shown in **Figure 6**.

**Supplementary Figure 6**. Taxonomic profiles of the rectal microbiomes, paired with the endocervical and vaginal microbiomes (data not shown), based on relative abundance at the species at baseline and follow-up for the three study cohort datasets. The participant ID# is shown at the top of the respective box that contains baseline (left) and follow-up (right) bar graphs and the in enterotype noted below the bar graphs.  The samples exhibiting genito-rectal transmissions are shown in **Figure 6**.

**Supplementary Figure 7**: Function- and pathway-level community analysis of paired vaginal, endocervical and rectal microbiomes of women who did not have a *C. trachomatis* infection and did not receive treatment. A. The Function-level analysis shows the function that had significantly different relative abundances between baseline and follow-up microbiomes for the rectum. No functions were significantly different for the endocervical or vaginal sites. B. The microbial contribution for each function and the associated pathways are shown for baseline and follow up time points. C. The Pathway-level community analysis shows those pathways for each anatomic site that have a significant difference in relative abundance between the baseline and follow-up microbiomes ($P<0.05$). Species represented in the graphs are color-coded as per the columns to the right.

**Supplementary Figure 8**. MAFFT alignment of *C. trachomatis rpl*V genes (encoding L22 protein) extracted from baseline and follow-up samples of the *Ct* persistence cohort and baseline samples of the *Ct* clearance group. *Ct* reference strain D/UW-3/CX was used as the sequence to identify the macrolide conferring resistance mutations in our clinical samples. Regions with nonsynonymous missense mutations are shown. Numbering above the nucleotide sequence indicates nucleotide position and numbering below the amino acid sequence indicates amino acid position. The macrolide binding site is noted in blue below the amino acid sequence. Presence of *rpl*V mutations in *Ct* is indicative of macrolide resistance as reported by Mestrovic et al., 2018 and Benamri et al., 2021.

**Supplementary Figure 9.** MAFFT alignment of *L. iners erm*B genes extracted from baseline and follow-up samples across all three cohorts, *L. iners* strain C0094A1 and C0210C1 harboring *erm*B and the ErmB protein homolog from *Enterococcus faecium* (available from CARD). Regions with nonsynonymous missense mutations are shown. Numbering above the nucleotide sequence indicates nucleotide position and numbering below the amino acid sequence indicates amino acid position. The *L. iners erm*B genes circulating in our microbiomes were found to be >/= 99.1% identical to the protein homolog model, and the strains C0094A1 and C0210C1 were 99.1% identical. Presence of *erm*B in *L. iners* is indicative of macrolide resistance as reported by the CARD model: https://card.mcmaster.ca/ontology/36514.

**Supplementary Figure 10.** MAFFT alignment of *G. vaginalis ermX* genes extracted from baseline and follow-up samples across all three cohorts, *G. vaginalis* strain JNFY11 harboring *erm*X and the ErmX protein homolog from plasmid pNG2 (available from CARD). Regions with nonsynonymous missense mutations are shown. Numbering above the nucleotide sequence indicates nucleotide position and numbering below the amino acid sequence indicates amino acid position. The. *G vaginalis erm*X genes circulating in our microbiomes were found to be >/= 93.8% identical to the protein homolog model and the strain JNFY11 was 99.3% identical. Presence of *erm*X in *G. vaginalis* is indicative of macrolide resistance as reported by the CARD model: https://card.mcmaster.ca/ontology/36735. Presence of more than one copy of *erm*X represented at _2 next to a given sample ID is indicative of the presence of a second copy of the *G. vaginalis erm*X gene within a microbiome.

**Supplementary Tables**

**Supplementary Table S1.** Metagenomic shotgun sequencing results and quality control statistics.

**Supplementary Table S2.** Metadata of samples used in this study, including data on age, ethnicity, C. trachomatis, BV, pH, *N. gonorrhoeae*, *M. genitalium* and HPV type.

**Supplementary Table S3.** Distribution of low risk (lr)HPV (black font) and high risk (hr)HPV (red & orange fonts) types in paired vaginal, endocervical and rectal microbiomes.

**Supplementary Table S4**. Comparison of subCommunity State Type (CST) and metagenomic (mg)CST in paired vaginal and endocervical microbiomes for the three cohort datasets at baseline.

**Supplementary Table S5**. Comparison of vaginal and endocervical subCST and mgCST classification systems and associations with BV and *C. trachomatis* (*Ct*).

**Supplementary Table S6**. subCST associations with BV and *C. trachomatis* infection in the vaginal and endocervical microbiomes.

**Supplementary Table S7**. Functions and pathways associated with the baseline time point for the *C. trachomatis* persistence cohort microbiomes.

**Supplementary Table S8**. Functions and pathways associated with the follow up time point for the *C. trachomatis* persistence cohort microbiomes.

**Supplementary Table S9**. Functions and pathways associated with the baseline time point for the *C. trachomatis* clearance cohort microbiomes.

**Supplementary Table S10**. Functions and pathways associated with the follow up time point for the *C. trachomatis* clearance cohort microbiomes.

**Supplementary Table S11**. Functions and pathways associated with the baseline time point for the *C. trachomatis* control cohort microbiomes.

**Supplementary Table S12**. Functions and pathways associated with the follow up time point for the *C. trachomatis* control cohort microbiomes.

**Supplementary Table S13**. Function-level community analysis of the microbiomes from the *Ct* clearance cohort.

**Supplementary Table S14**. Function-level community analysis of the microbiomes from the *Ct* persistence cohort.

**Supplementary Table S15**. Function-level community analysis of the microbiomes from the no treatment cohort.

**Supplementary Table S16**. *G. vaginalis erm*X, and *L. iners erm*B and *tet*(W) genes and mutations in the *C. trachomatis rpl*V gene conferring resistance to macrolide antibiotics identified among rectal, vaginal and endocervical microbiomes for all three cohorts at baseline (blue) and follow-up (red).

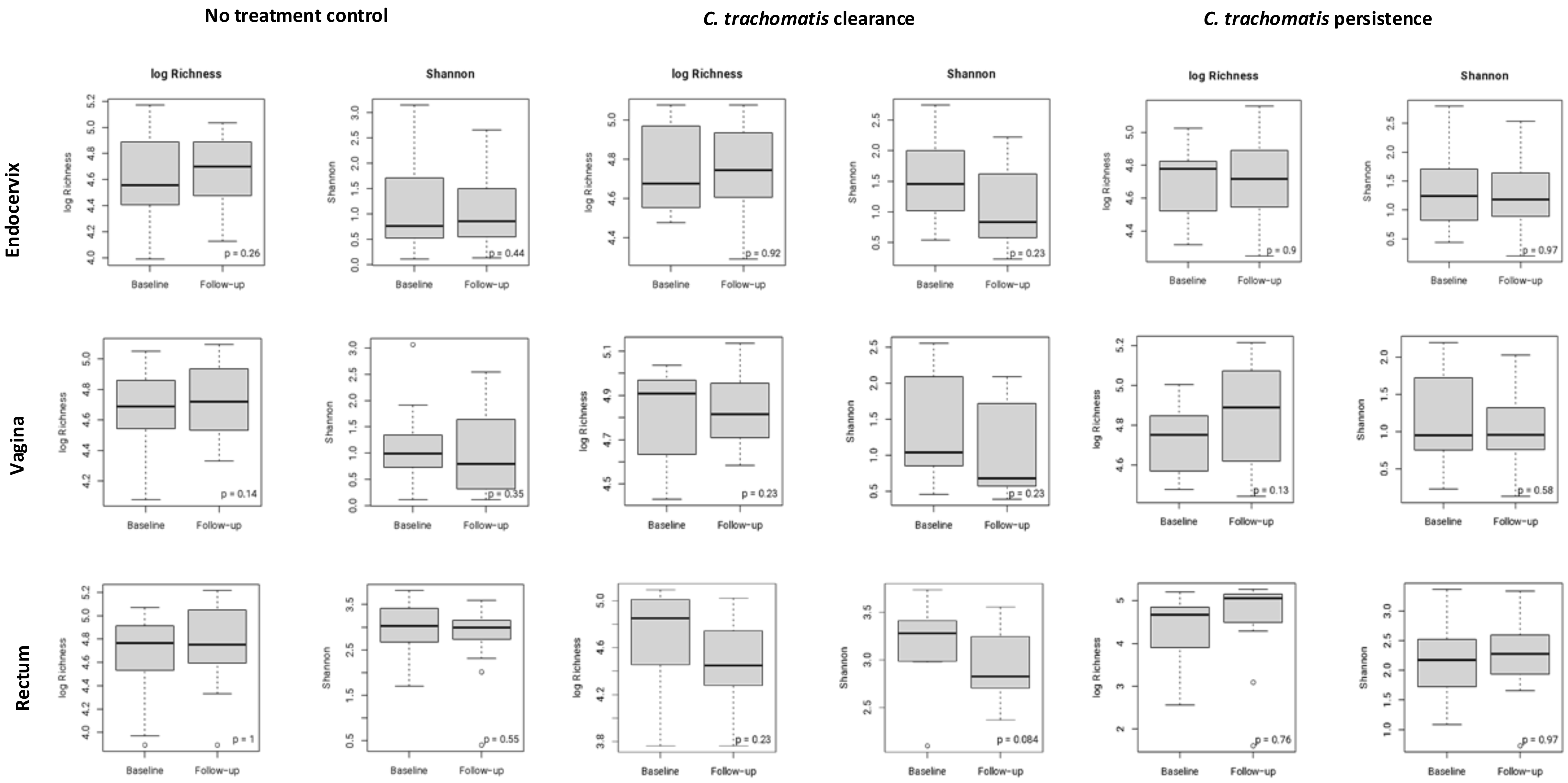
No treatment control
C. trachomatis clearance
C. trachomatis persistence
Endocervix
Vagina
Rectum
log Richness
Shannon
Baseline
Follow-up
p = 0.26
p = 0.44
p = 0.92
p = 0.23
p = 0.9
p = 0.97
p = 0.14
p = 0.35
p = 0.23
p = 0.23
p = 0.13
p = 0.58
p = 1
p = 0.55
p = 0.23
p = 0.084
p = 0.76
p = 0.97

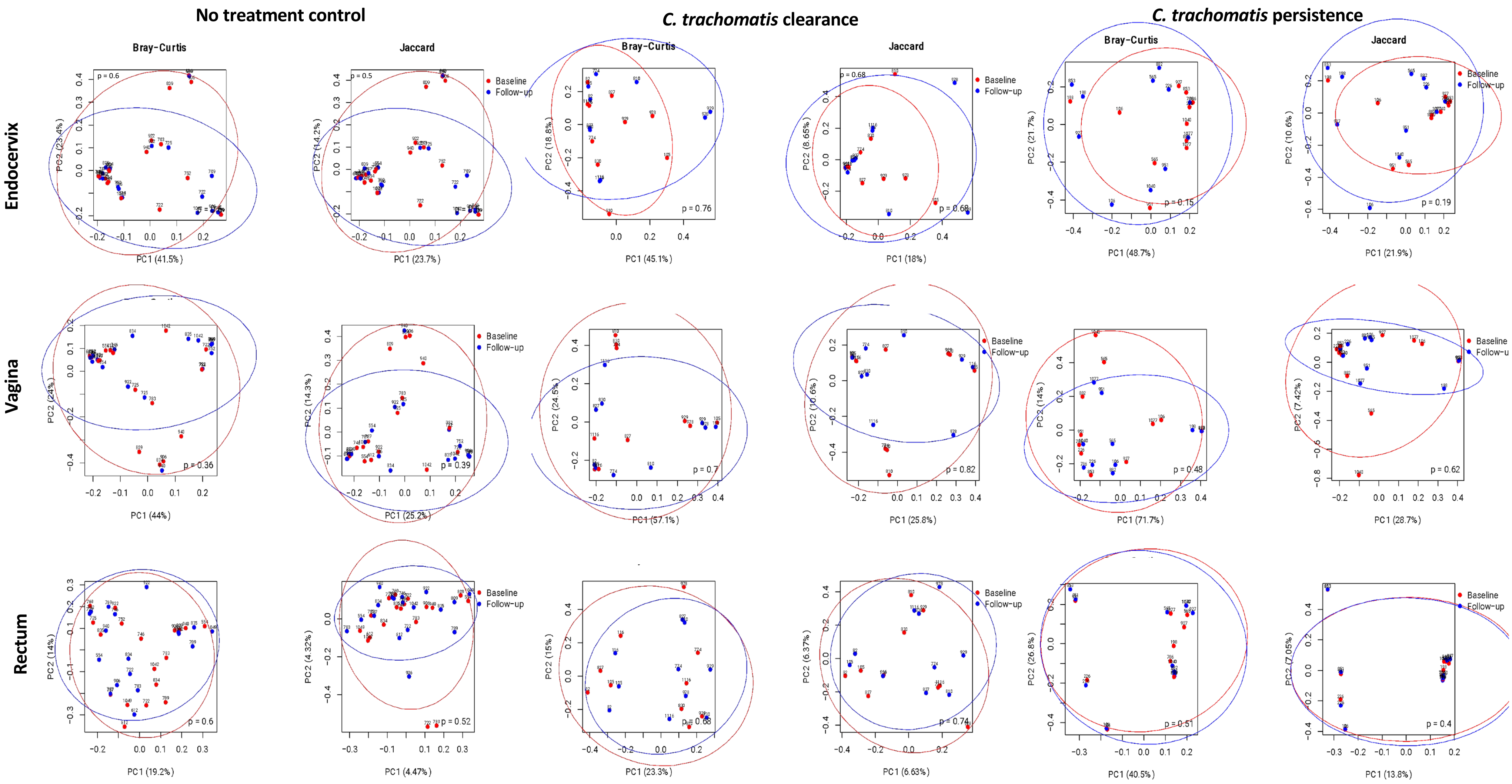
No treatment control
C. trachomatis clearance
C. trachomatis persistence
Bray-Curtis
Jaccard
Endocervix
Vagina
Rectum
Baseline
Follow-up
p = 0.6
PC1 (41.5%)
PC2 (23.4%)
p = 0.5
PC1 (23.7%)
PC2 (14.2%)
p = 0.76
PC1 (45.1%)
PC2 (18.8%)
p = 0.68
PC1 (18%)
PC2 (8.65%)
p = 0.15
PC1 (48.7%)
PC2 (21.7%)
p = 0.19
PC1 (21.9%)
PC2 (10.6%)
p = 0.36
PC1 (44%)
PC2 (24%)
p = 0.39
PC1 (25.2%)
PC2 (14.3%)
p = 0.7
PC1 (57.1%)
PC2 (24.5%)
p = 0.82
PC1 (25.8%)
PC2 (10.6%)
p = 0.48
PC1 (71.7%)
PC2 (14%)
p = 0.62
PC1 (28.7%)
PC2 (7.42%)
p = 0.6
PC1 (19.2%)
PC2 (14%)
p = 0.52
PC1 (4.47%)
PC2 (4.32%)
p = 0.68
PC1 (23.3%)
PC2 (15%)
p = 0.74
PC1 (6.63%)
PC2 (6.37%)
p = 0.51
PC1 (40.5%)
PC2 (26.8%)
p = 0.4
PC1 (13.8%)
PC2 (7.05%)

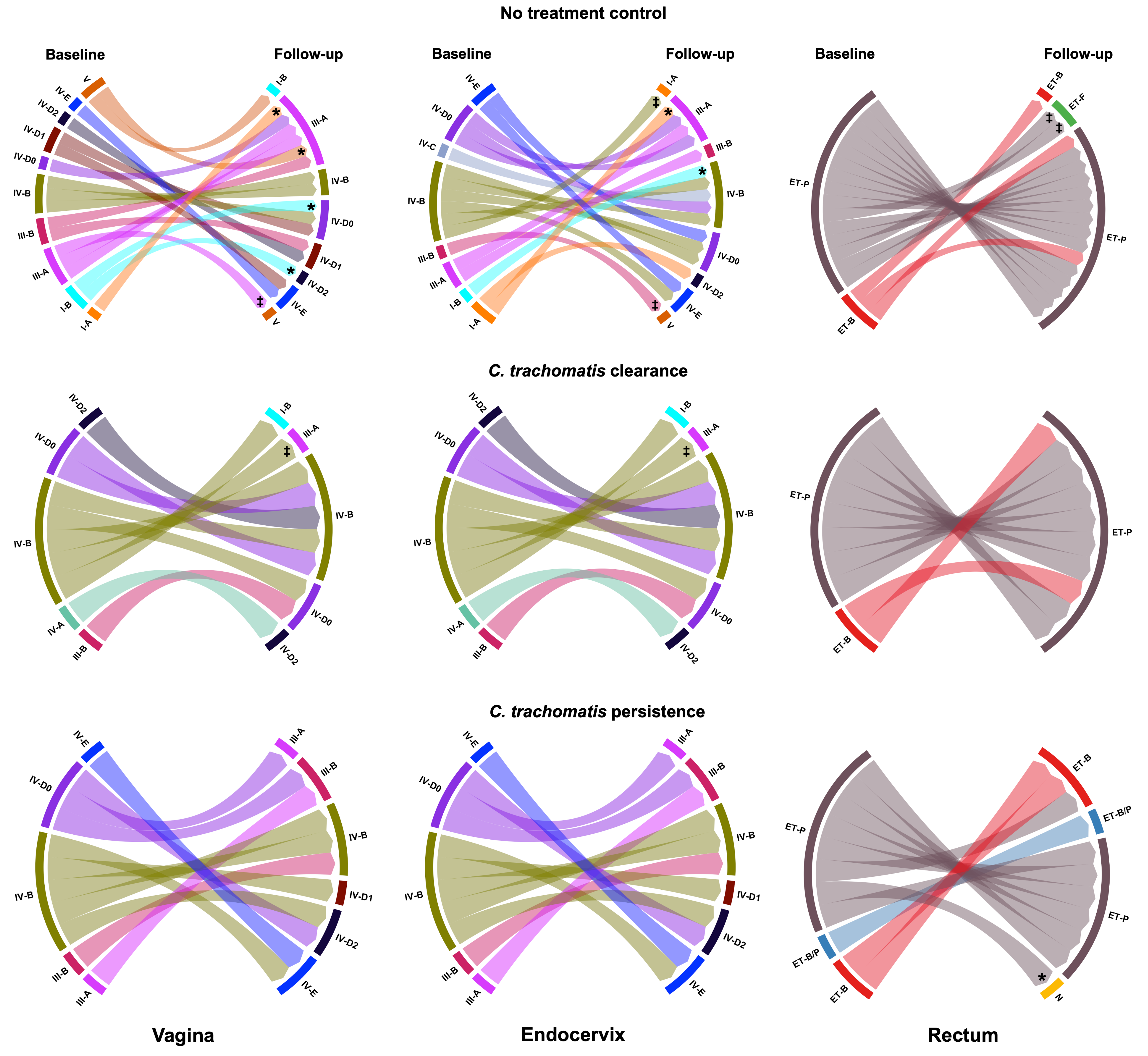
No treatment control
Baseline
Follow-up
V
IV-E
IV-D2
IV-D1
IV-D0
IV-B
III-B
III-A
I-B
I-A
III-A
IV-B
IV-D0
IV-D1
IV-D2
IV-E
V
*
‡
IV-C
III-B
ET-B
ET-F
ET-P
C. trachomatis clearance
IV-A
C. trachomatis persistence
ET-B/P
N
Vagina
Endocervix
Rectum

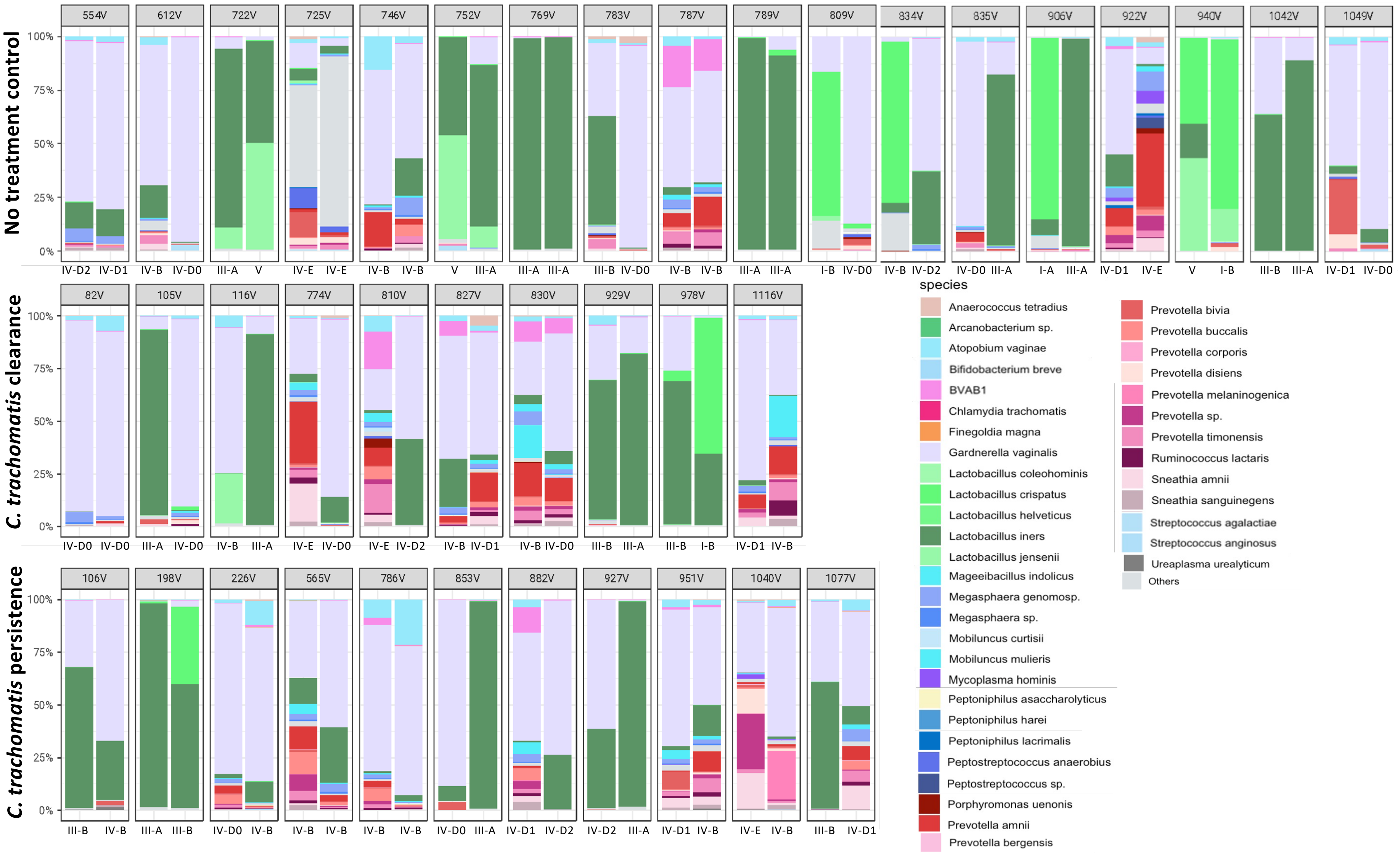

No treatment control
C. trachomatis clearance
C. trachomatis persistence
100%
75%
50%
25%
0%
554V
612V
722V
725V
746V
752V
769V
783V
787V
789V
809V
834V
835V
906V
922V
940V
1042V
1049V
IV-D2 IV-D1
IV-B IV-D0
III-A V
IV-E IV-E
IV-B IV-B
V III-A
III-A III-A
III-B IV-D0
IV-B IV-B
III-A III-A
I-B IV-D0
IV-B IV-D2
IV-D0 III-A
I-A III-A
IV-D1 IV-E
V I-B
III-B III-A
IV-D1 IV-D0
82V
105V
116V
774V
810V
827V
830V
929V
978V
1116V
IV-D0 IV-D0
III-A IV-D0
IV-B III-A
IV-E IV-D0
IV-E IV-D2
IV-B IV-D1
IV-B IV-D0
III-B III-A
III-B I-B
IV-D1 IV-B
106V
198V
226V
565V
786V
853V
882V
927V
951V
1040V
1077V
III-B IV-B
III-A III-B
IV-D0 IV-B
IV-B IV-B
IV-B IV-B
IV-D0 III-A
IV-D1 IV-D2
IV-D2 III-A
IV-D1 IV-B
IV-E IV-B
III-B IV-D1
species
Anaerococcus tetradius
Arcanobacterium sp.
Atopobium vaginae
Bifidobacterium breve
BVAB1
Chlamydia trachomatis
Finegoldia magna
Gardnerella vaginalis
Lactobacillus coleohominis
Lactobacillus crispatus
Lactobacillus helveticus
Lactobacillus iners
Lactobacillus jensenii
Mageeibacillus indolicus
Megasphaera genomosp.
Megasphaera sp.
Mobiluncus curtisii
Mobiluncus mulieris
Mycoplasma hominis
Peptoniphilus asaccharolyticus
Peptoniphilus harei
Peptoniphilus lacrimalis
Peptostreptococcus anaerobius
Peptostreptococcus sp.
Porphyromonas uenonis
Prevotella amnii
Prevotella bergensis
Prevotella bivia
Prevotella buccalis
Prevotella corporis
Prevotella disiens
Prevotella melaninogenica
Prevotella sp.
Prevotella timonensis
Ruminococcus lactaris
Sneathia amnii
Sneathia sanguinegens
Streptococcus agalactiae
Streptococcus anginosus
Ureaplasma urealyticum
Others

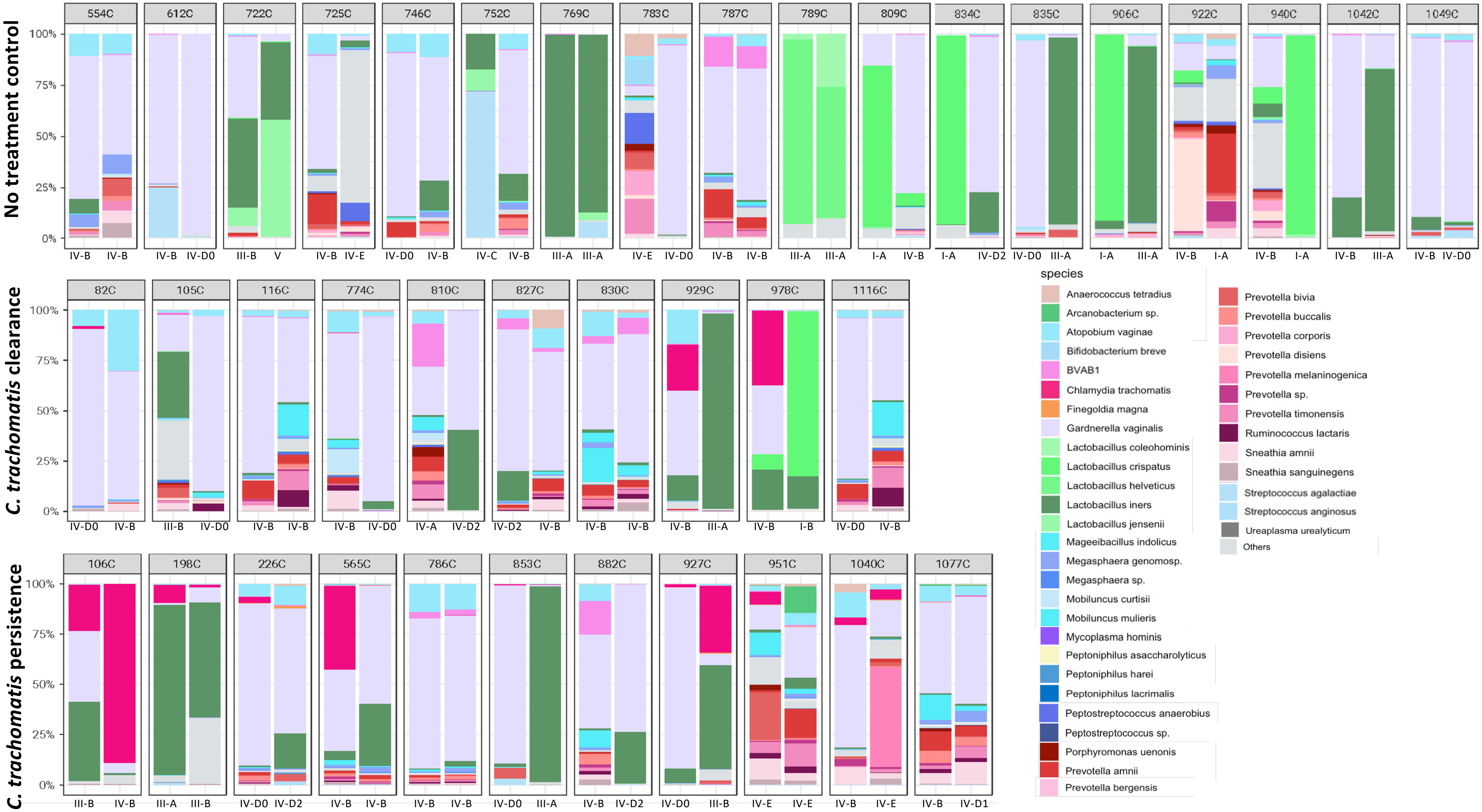

No treatment control
554C
612C
722C
725C
746C
752C
769C
783C
787C
789C
809C
834C
835C
906C
922C
940C
1042C
1049C
100%
75%
50%
25%
0%
IV-B IV-B
IV-B IV-D0
III-B V
IV-B IV-E
IV-D0 IV-B
IV-C IV-B
III-A III-A
IV-E IV-D0
IV-B IV-B
III-A III-A
I-A IV-B
I-A IV-D2
IV-D0 III-A
I-A III-A
IV-B I-A
IV-B I-A
IV-B III-A
IV-B IV-D0
C. trachomatis clearance
82C
105C
116C
774C
810C
827C
830C
929C
978C
1116C
IV-D0 IV-B
III-B IV-D0
IV-B IV-B
IV-B IV-D0
IV-A IV-D2
IV-D2 IV-B
IV-B IV-B
IV-B III-A
IV-B I-B
IV-D0 IV-B
C. trachomatis persistence
106C
198C
226C
565C
786C
853C
882C
927C
951C
1040C
1077C
III-B IV-B
III-A III-B
IV-D0 IV-D2
IV-B IV-B
IV-B IV-B
IV-D0 III-A
IV-B IV-D2
IV-D0 III-B
IV-E IV-E
IV-B IV-E
IV-B IV-D1
species
Anaerococcus tetradius
Arcanobacterium sp.
Atopobium vaginae
Bifidobacterium breve
BVAB1
Chlamydia trachomatis
Finegoldia magna
Gardnerella vaginalis
Lactobacillus coleohominis
Lactobacillus crispatus
Lactobacillus helveticus
Lactobacillus iners
Lactobacillus jensenii
Mageeibacillus indolicus
Megasphaera genomosp.
Megasphaera sp.
Mobiluncus curtisii
Mobiluncus mulieris
Mycoplasma hominis
Peptoniphilus asaccharolyticus
Peptoniphilus harei
Peptoniphilus lacrimalis
Peptostreptococcus anaerobius
Peptostreptococcus sp.
Porphyromonas uenonis
Prevotella amnii
Prevotella bergensis
Prevotella bivia
Prevotella buccalis
Prevotella corporis
Prevotella disiens
Prevotella melaninogenica
Prevotella sp.
Prevotella timonensis
Ruminococcus lactaris
Sneathia amnii
Sneathia sanguinegens
Streptococcus agalactiae
Streptococcus anginosus
Ureaplasma urealyticum
Others

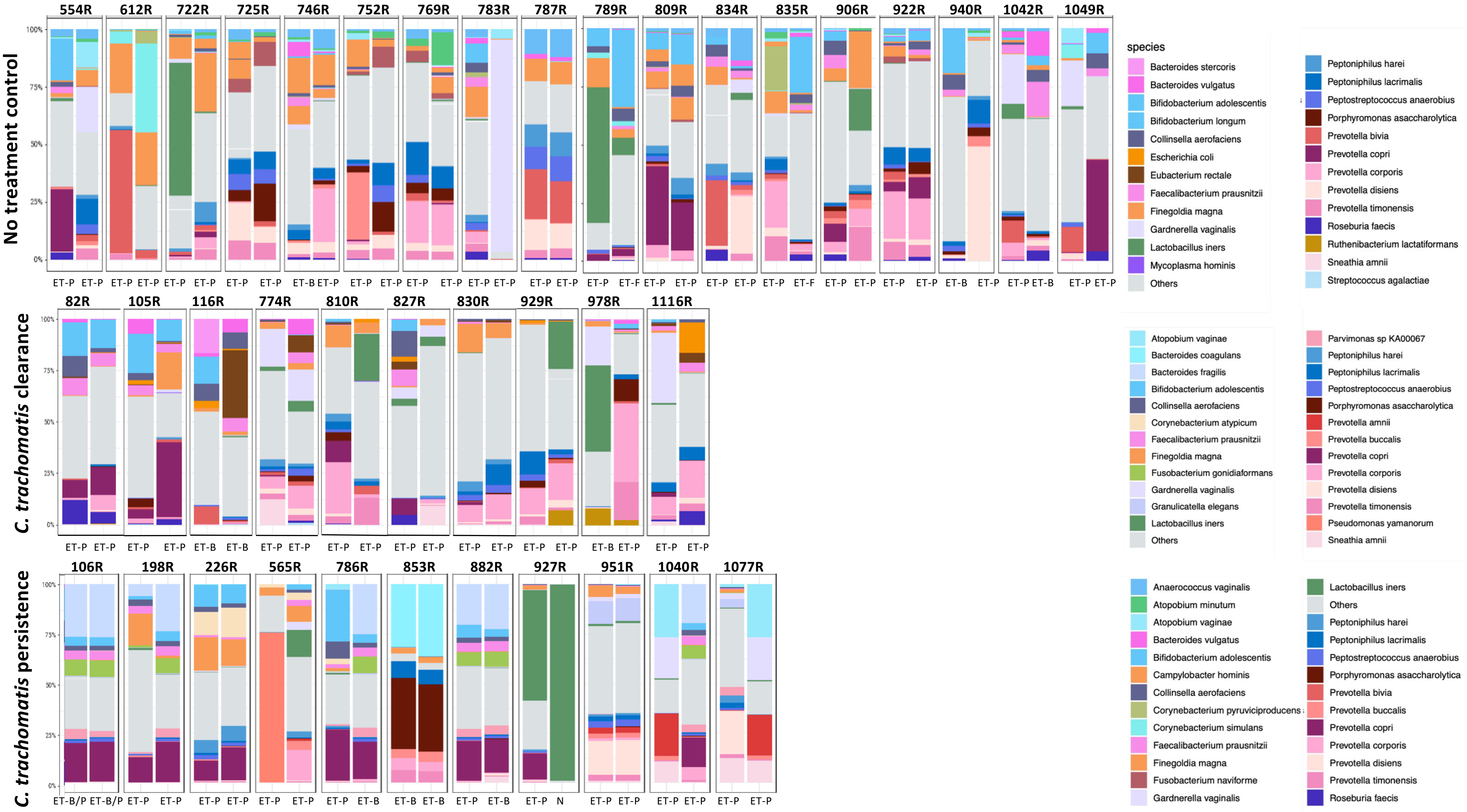

No treatment control
554R
612R
722R
725R
746R
752R
769R
783R
787R
789R
809R
834R
835R
906R
922R
940R
1042R
1049R
C. trachomatis clearance
82R
105R
116R
774R
810R
827R
830R
929R
978R
1116R
C. trachomatis persistence
106R
198R
226R
565R
786R
853R
882R
927R
951R
1040R
1077R
100%
75%
50%
25%
0%
ET-P
ET-B
ET-F
ET-B/P
N
species
Bacteroides stercoris
Bacteroides vulgatus
Bifidobacterium adolescentis
Bifidobacterium longum
Collinsella aerofaciens
Escherichia coli
Eubacterium rectale
Faecalibacterium prausnitzii
Finegoldia magna
Gardnerella vaginalis
Lactobacillus iners
Mycoplasma hominis
Others
Peptoniphilus harei
Peptoniphilus lacrimalis
Peptostreptococcus anaerobius
Porphyromonas asaccharolytica
Prevotella bivia
Prevotella copri
Prevotella corporis
Prevotella disiens
Prevotella timonensis
Roseburia faecis
Ruthenibacterium lactatiformans
Sneathia amnii
Streptococcus agalactiae
Atopobium vaginae
Bacteroides coagulans
Bacteroides fragilis
Corynebacterium atypicum
Fusobacterium gonidiaformans
Granulicatella elegans
Parvimonas sp KA00067
Prevotella amnii
Prevotella buccalis
Pseudomonas yamanorum
Anaerococcus vaginalis
Atopobium minutum
Campylobacter hominis
Corynebacterium pyruviciproducens
Corynebacterium simulans
Fusobacterium naviforme

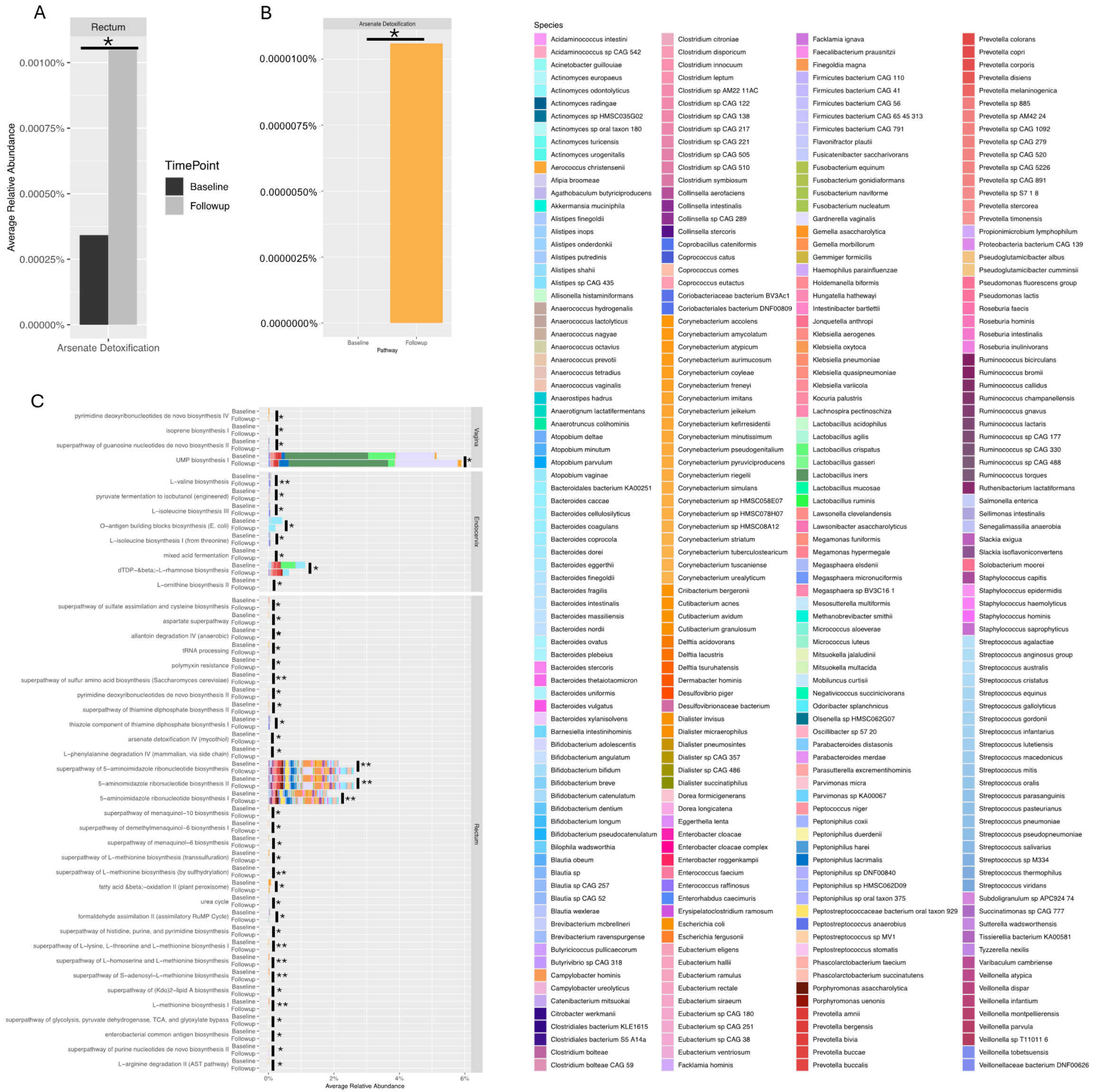

A
Rectum
*
0.00100%
0.00075%
0.00050%
0.00025%
0.00000%
Average Relative Abundance
Arsenate Detoxification
TimePoint
Baseline
Followup
B
Arsenate Detoxification
*
0.0000100%
0.0000075%
0.0000050%
0.0000025%
0.0000000%
Baseline
Followup
Pathway
C
pyrimidine deoxyribonucleotides de novo biosynthesis IV
isoprene biosynthesis I
superpathway of guanosine nucleotides de novo biosynthesis II
UMP biosynthesis I
L−valine biosynthesis
pyruvate fermentation to isobutanol (engineered)
L−isoleucine biosynthesis III
O−antigen building blocks biosynthesis (E. coli)
L−isoleucine biosynthesis I (from threonine)
mixed acid fermentation
dTDP−β−L−rhamnose biosynthesis
L−ornithine biosynthesis II
superpathway of sulfate assimilation and cysteine biosynthesis
aspartate superpathway
allantoin degradation IV (anaerobic)
tRNA processing
polymyxin resistance
superpathway of sulfur amino acid biosynthesis (Saccharomyces cerevisiae)
pyrimidine deoxyribonucleotides de novo biosynthesis II
superpathway of thiamine diphosphate biosynthesis II
thiazole component of thiamine diphosphate biosynthesis I
arsenate detoxification IV (mycothiol)
L−phenylalanine degradation IV (mammalian, via side chain)
superpathway of 5−aminoimidazole ribonucleotide biosynthesis
5−aminoimidazole ribonucleotide biosynthesis II
5−aminoimidazole ribonucleotide biosynthesis I
superpathway of menaquinol−10 biosynthesis
superpathway of demethylmenaquinol−6 biosynthesis I
superpathway of menaquinol−6 biosynthesis
superpathway of L−methionine biosynthesis (transsulfuration)
superpathway of L−methionine biosynthesis (by sulfhydrylation)
fatty acid β−oxidation II (plant peroxisome)
urea cycle
formaldehyde assimilation II (assimilatory RuMP Cycle)
superpathway of histidine, purine, and pyrimidine biosynthesis
superpathway of L−lysine, L−threonine and L−methionine biosynthesis I
superpathway of L−homoserine and L−methionine biosynthesis
superpathway of S−adenosyl−L−methionine biosynthesis
superpathway of (Kdo)2−lipid A biosynthesis
L−methionine biosynthesis I
superpathway of glycolysis, pyruvate dehydrogenase, TCA, and glyoxylate bypass
enterobacterial common antigen biosynthesis
superpathway of purine nucleotides de novo biosynthesis II
L−arginine degradation II (AST pathway)
Baseline
Followup
*
**
Vagina
Endocervix
Rectum
0%
2%
4%
6%
Average Relative Abundance
Species
Acidaminococcus intestini
Acidaminococcus sp CAG 542
Acinetobacter guillouiae
Actinomyces europaeus
Actinomyces odontolyticus
Actinomyces radingae
Actinomyces sp HMSC035G02
Actinomyces sp oral taxon 180
Actinomyces turicensis
Actinomyces urogenitalis
Aerococcus christensenii
Afipia broomeae
Agathobaculum butyriciproducens
Akkermansia muciniphila
Alistipes finegoldii
Alistipes inops
Alistipes onderdonkii
Alistipes putredinis
Alistipes shahii
Alistipes sp CAG 435
Allisonella histaminiformans
Anaerococcus hydrogenalis
Anaerococcus lactolyticus
Anaerococcus nagyae
Anaerococcus octavius
Anaerococcus prevotii
Anaerococcus tetradius
Anaerococcus vaginalis
Anaerostipes hadrus
Anaerotignum lactatifermentans
Anaerotruncus colihominis
Atopobium deltae
Atopobium minutum
Atopobium parvulum
Atopobium vaginae
Bacteroidales bacterium KA00251
Bacteroides caccae
Bacteroides cellulosilyticus
Bacteroides coagulans
Bacteroides coprocola
Bacteroides dorei
Bacteroides eggerthii
Bacteroides finegoldii
Bacteroides fragilis
Bacteroides intestinalis
Bacteroides massiliensis
Bacteroides nordii
Bacteroides ovatus
Bacteroides plebeius
Bacteroides stercoris
Bacteroides thetaiotaomicron
Bacteroides uniformis
Bacteroides vulgatus
Bacteroides xylanisolvens
Barnesiella intestinihominis
Bifidobacterium adolescentis
Bifidobacterium angulatum
Bifidobacterium bifidum
Bifidobacterium breve
Bifidobacterium catenulatum
Bifidobacterium dentium
Bifidobacterium longum
Bifidobacterium pseudocatenulatum
Bilophila wadsworthia
Blautia obeum
Blautia sp
Blautia sp CAG 257
Blautia sp CAG 52
Blautia wexlerae
Brevibacterium mcbrellneri
Brevibacterium ravenspurgense
Butyricicoccus pullicaecorum
Butyrivibrio sp CAG 318
Campylobacter hominis
Campylobacter ureolyticus
Catenibacterium mitsuokai
Citrobacter werkmanii
Clostridiales bacterium KLE1615
Clostridiales bacterium S5 A14a
Clostridium bolteae
Clostridium bolteae CAG 59
Clostridium citroniae
Clostridium disporicum
Clostridium innocuum
Clostridium leptum
Clostridium sp AM22 11AC
Clostridium sp CAG 122
Clostridium sp CAG 138
Clostridium sp CAG 217
Clostridium sp CAG 221
Clostridium sp CAG 505
Clostridium sp CAG 510
Clostridium symbiosum
Collinsella aerofaciens
Collinsella intestinalis
Collinsella sp CAG 289
Collinsella stercoris
Coprobacillus cateniformis
Coprococcus catus
Coprococcus comes
Coprococcus eutactus
Coriobacteriaceae bacterium BV3Ac1
Coriobacteriales bacterium DNF00809
Corynebacterium accolens
Corynebacterium amycolatum
Corynebacterium atypicum
Corynebacterium aurimucosum
Corynebacterium coyleae
Corynebacterium freneyi
Corynebacterium imitans
Corynebacterium jeikeium
Corynebacterium kefirresidentii
Corynebacterium minutissimum
Corynebacterium pseudogenitalium
Corynebacterium pyruviciproducens
Corynebacterium riegelii
Corynebacterium simulans
Corynebacterium sp HMSC058E07
Corynebacterium sp HMSC078H07
Corynebacterium sp HMSC08A12
Corynebacterium striatum
Corynebacterium tuberculostearicum
Corynebacterium tuscaniense
Corynebacterium urealyticum
Criibacterium bergeronii
Cutibacterium acnes
Cutibacterium avidum
Cutibacterium granulosum
Delftia acidovorans
Delftia lacustris
Delftia tsuruhatensis
Dermabacter hominis
Desulfovibrio piger
Desulfovibrionaceae bacterium
Dialister invisus
Dialister micraerophilus
Dialister pneumosintes
Dialister sp CAG 357
Dialister sp CAG 486
Dialister succinatiphilus
Dorea formicigenerans
Dorea longicatena
Eggerthella lenta
Enterobacter cloacae
Enterobacter cloacae complex
Enterobacter roggenkampii
Enterococcus faecium
Enterococcus raffinosus
Enterorhabdus caecimuris
Erysipelatoclostridium ramosum
Escherichia coli
Escherichia fergusonii
Eubacterium eligens
Eubacterium hallii
Eubacterium ramulus
Eubacterium rectale
Eubacterium siraeum
Eubacterium sp CAG 180
Eubacterium sp CAG 251
Eubacterium sp CAG 38
Eubacterium ventriosum
Facklamia hominis
Facklamia ignava
Faecalibacterium prausnitzii
Finegoldia magna
Firmicutes bacterium CAG 110
Firmicutes bacterium CAG 41
Firmicutes bacterium CAG 56
Firmicutes bacterium CAG 65 45 313
Firmicutes bacterium CAG 791
Flavonifractor plautii
Fusicatenibacter saccharivorans
Fusobacterium equinum
Fusobacterium gonidiaformans
Fusobacterium naviforme
Fusobacterium nucleatum
Gardnerella vaginalis
Gemella asaccharolytica
Gemella morbillorum
Gemmiger formicilis
Haemophilus parainfluenzae
Holdemanella biformis
Hungatella hathewayi
Intestinibacter bartlettii
Jonquetella anthropi
Klebsiella aerogenes
Klebsiella oxytoca
Klebsiella pneumoniae
Klebsiella quasipneumoniae
Klebsiella variicola
Kocuria palustris
Lachnospira pectinoschiza
Lactobacillus acidophilus
Lactobacillus agilis
Lactobacillus crispatus
Lactobacillus gasseri
Lactobacillus iners
Lactobacillus mucosae
Lactobacillus ruminis
Lawsonella clevelandensis
Lawsonibacter asaccharolyticus
Megamonas funiformis
Megamonas hypermegale
Megasphaera elsdenii
Megasphaera micronuciformis
Megasphaera sp BV3C16 1
Mesosutterella multiformis
Methanobrevibacter smithii
Micrococcus aloeverae
Micrococcus luteus
Mitsuokella jalaludinii
Mitsuokella multacida
Mobiluncus curtisii
Negativicoccus succinicivorans
Odoribacter splanchnicus
Olsenella sp HMSC062G07
Oscillibacter sp 57 20
Parabacteroides distasonis
Parabacteroides merdae
Parasutterella excrementihominis
Parvimonas micra
Parvimonas sp KA00067
Peptococcus niger
Peptoniphilus coxii
Peptoniphilus duerdenii
Peptoniphilus harei
Peptoniphilus lacrimalis
Peptoniphilus sp DNF00840
Peptoniphilus sp HMSC062D09
Peptoniphilus sp oral taxon 375
Peptostreptococcaceae bacterium oral taxon 929
Peptostreptococcus anaerobius
Peptostreptococcus sp MV1
Peptostreptococcus stomatis
Phascolarctobacterium faecium
Phascolarctobacterium succinatutens
Porphyromonas asaccharolytica
Porphyromonas uenonis
Prevotella amnii
Prevotella bergensis
Prevotella bivia
Prevotella buccae
Prevotella buccalis
Prevotella colorans
Prevotella copri
Prevotella corporis
Prevotella disiens
Prevotella melaninogenica
Prevotella sp 885
Prevotella sp AM42 24
Prevotella sp CAG 1092
Prevotella sp CAG 279
Prevotella sp CAG 520
Prevotella sp CAG 5226
Prevotella sp CAG 891
Prevotella sp S7 1 8
Prevotella stercorea
Prevotella timonensis
Propionimicrobium lymphophilum
Proteobacteria bacterium CAG 139
Pseudoglutamicibacter albus
Pseudoglutamicibacter cumminsii
Pseudomonas fluorescens group
Pseudomonas lactis
Roseburia faecis
Roseburia hominis
Roseburia intestinalis
Roseburia inulinivorans
Ruminococcus bicirculans
Ruminococcus bromii
Ruminococcus callidus
Ruminococcus champanellensis
Ruminococcus gnavus
Ruminococcus lactaris
Ruminococcus sp CAG 177
Ruminococcus sp CAG 330
Ruminococcus sp CAG 488
Ruminococcus torques
Ruthenibacterium lactatiformans
Salmonella enterica
Sellimonas intestinalis
Senegalimassilia anaerobia
Slackia exigua
Slackia isoflavoniconvertens
Solobacterium moorei
Staphylococcus capitis
Staphylococcus epidermidis
Staphylococcus haemolyticus
Staphylococcus hominis
Staphylococcus saprophyticus
Streptococcus agalactiae
Streptococcus anginosus group
Streptococcus australis
Streptococcus cristatus
Streptococcus equinus
Streptococcus gallolyticus
Streptococcus gordonii
Streptococcus infantarius
Streptococcus lutetiensis
Streptococcus macedonicus
Streptococcus mitis
Streptococcus oralis
Streptococcus parasanguinis
Streptococcus pasteurianus
Streptococcus pneumoniae
Streptococcus pseudopneumoniae
Streptococcus salivarius
Streptococcus sp M334
Streptococcus thermophilus
Streptococcus viridans
Subdoligranulum sp APC924 74
Succinatimonas sp CAG 777
Sutterella wadsworthensis
Tissierella bacterium KA00581
Tyzzerella nexilis
Varibaculum cambriense
Veillonella atypica
Veillonella dispar
Veillonella infantium
Veillonella montpellierensis
Veillonella parvula
Veillonella sp T11011 6
Veillonella tobetsuensis
Veillonellaceae bacterium DNF00626

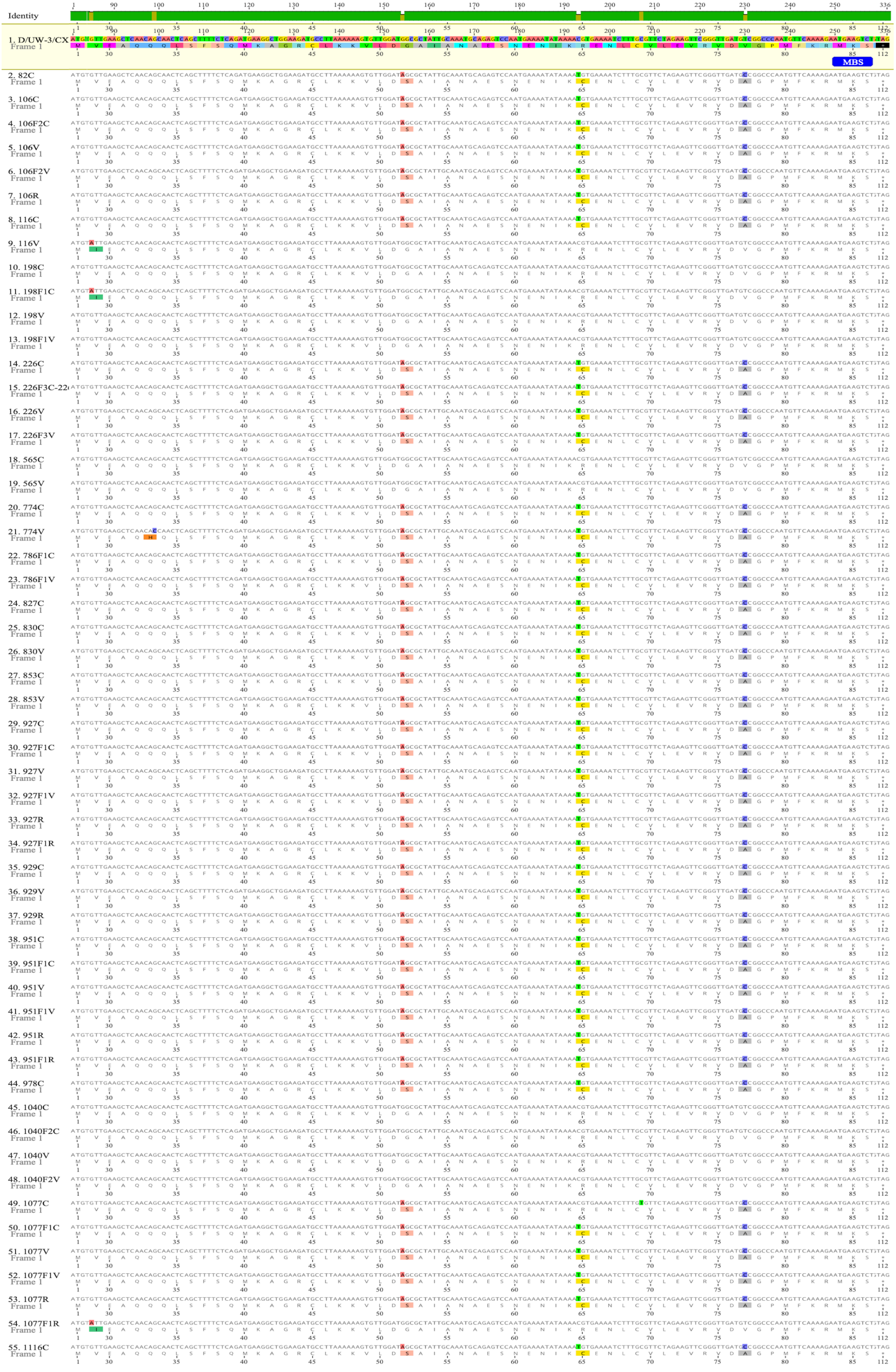
Identity
1. D/UW-3/CX
MBS
2. 82C
3. 106C
4. 106F2C
5. 106V
6. 106F2V
7. 106R
8. 116C
9. 116V
10. 198C
11. 198F1C
12. 198V
13. 198F1V
14. 226C
15. 226F3C-22
16. 226V
17. 226F3V
18. 565C
19. 565V
20. 774C
21. 774V
22. 786F1C
23. 786F1V
24. 827C
25. 830C
26. 830V
27. 853C
28. 853V
29. 927C
30. 927F1C
31. 927V
32. 927F1V
33. 927R
34. 927F1R
35. 929C
36. 929V
37. 929R
38. 951C
39. 951F1C
40. 951V
41. 951F1V
42. 951R
43. 951F1R
44. 978C
45. 1040C
46. 1040F2C
47. 1040V
48. 1040F2V
49. 1077C
50. 1077F1C
51. 1077V
52. 1077F1V
53. 1077R
54. 1077F1R
55. 1116C
Frame 1

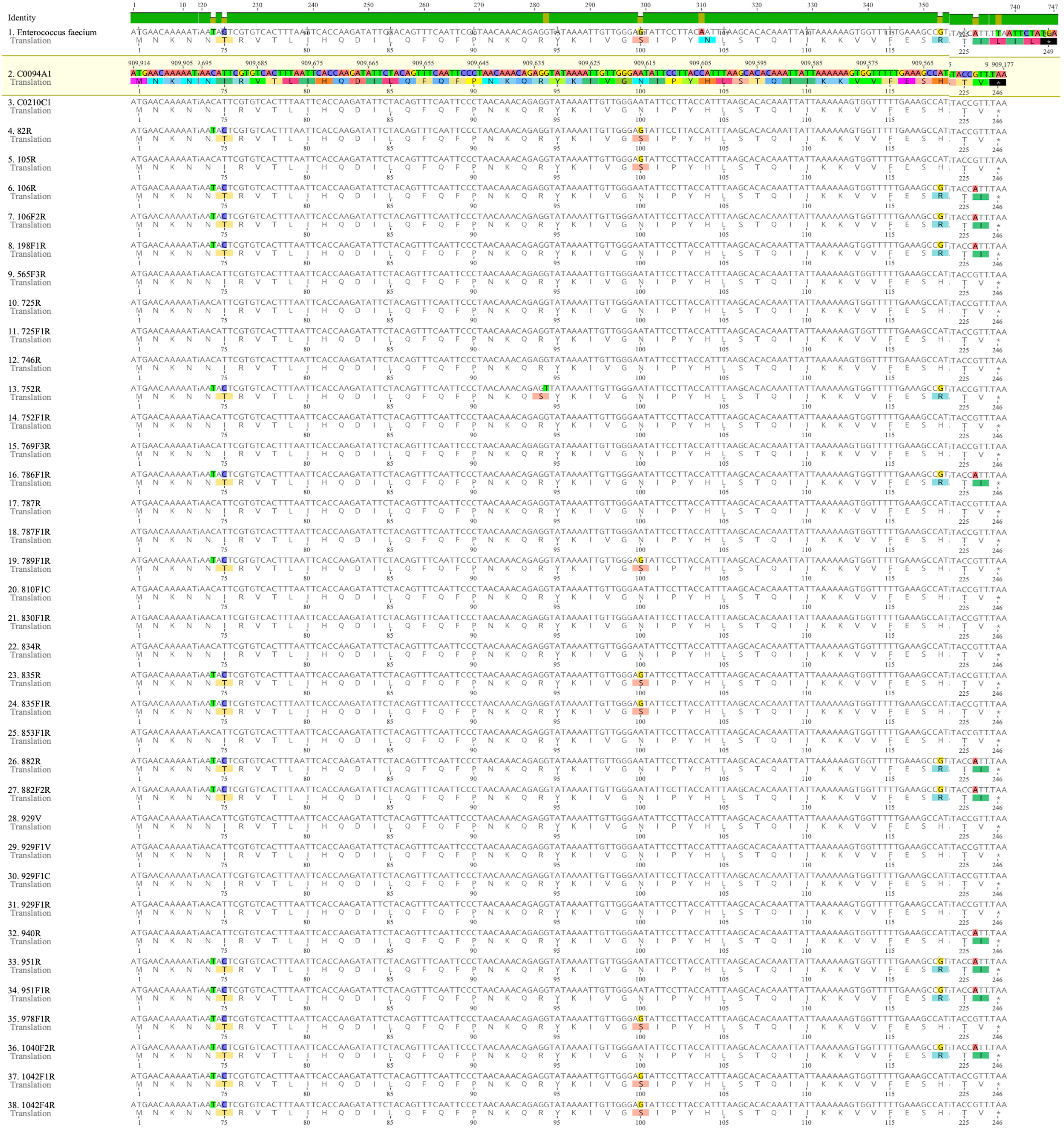
Identity
1. Enterococcus faecium
2. C0094A1
3. C0210C1
4. 82R
5. 105R
6. 106R
7. 106F2R
8. 198F1R
9. 565F3R
10. 725R
11. 725F1R
12. 746R
13. 752R
14. 752F1R
15. 769F3R
16. 786F1R
17. 787R
18. 787F1R
19. 789F1R
20. 810F1C
21. 830F1R
22. 834R
23. 835R
24. 835F1R
25. 853F1R
26. 882R
27. 882F2R
28. 929V
29. 929F1V
30. 929F1C
31. 929F1R
32. 940R
33. 951R
34. 951F1R
35. 978F1R
36. 1040F2R
37. 1042F1R
38. 1042F4R
Translation

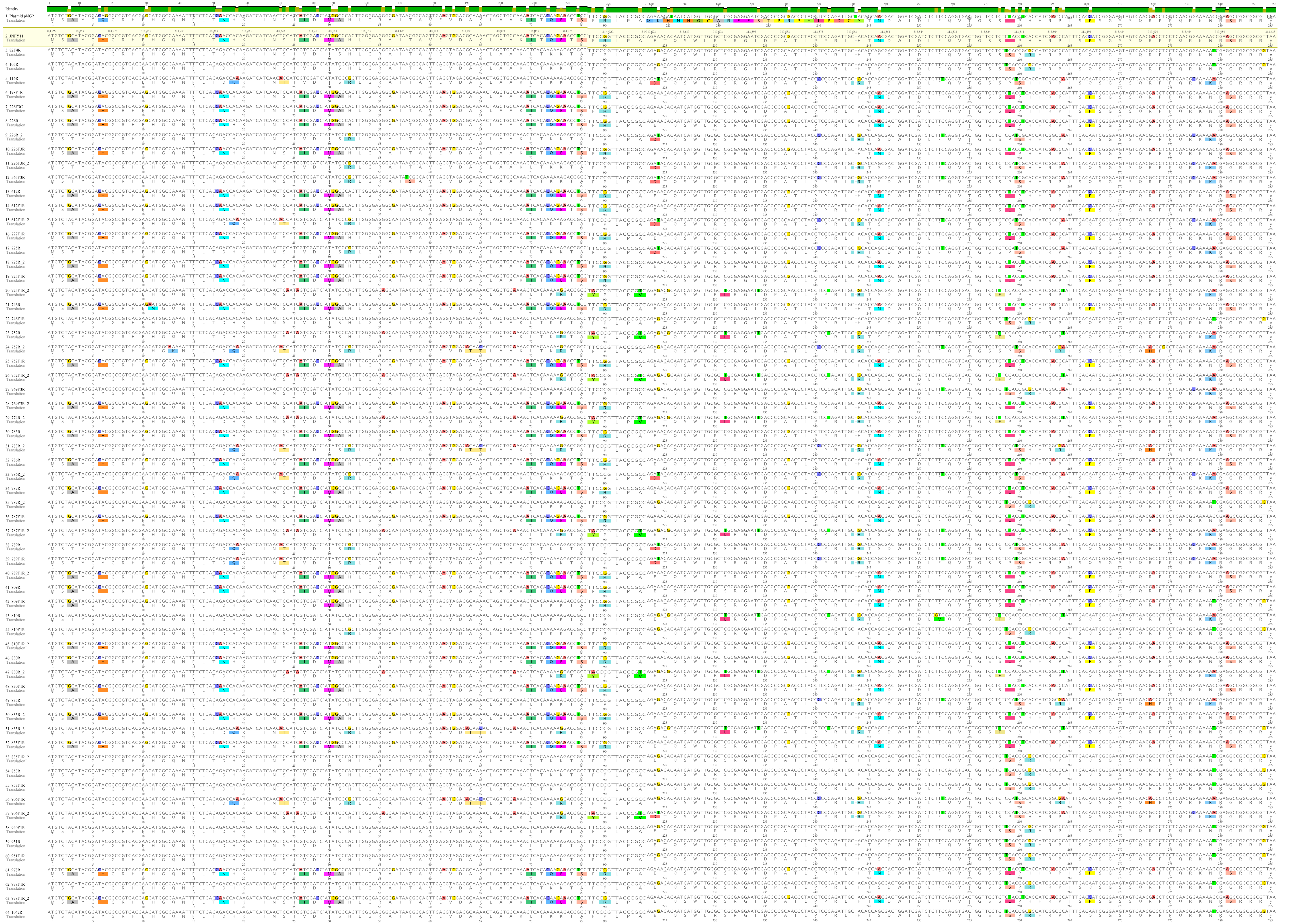
Identity
1. Plasmid pNG2
2. INFY11
3. 82F4R
4. 105R
5. 116R
6. 198F1R
7. 226F3C
8. 226R
9. 226R_2
10. 226F3R
11. 226F3R_2
12. 565F3R
13. 612R
14. 612F1R
15. 612F1R_2
16. 722F1R
17. 725R
18. 725R_2
19. 725F1R
20. 725F1R_2
21. 746R
22. 746F1R
23. 752R
24. 752R_2
25. 752F1R
26. 752F1R_2
27. 769F3R
28. 769F3R_2
29. 774R_2
30. 783R
31. 783R_2
32. 786R
33. 786R_2
34. 787R
35. 787R_2
36. 787F1R
37. 787F1R_2
38. 789R
39. 789F1R
40. 789F1R_2
41. 809R
42. 809F1R
43. 810R
44. 810F1R
45. 810F1R_2
46. 830R
47. 830R_2
48. 830F1R
49. 835R
50. 835R_2
51. 835R_3
52. 835F1R
53. 835F1R_2
54. 853R
55. 853F1R
56. 906F1R
57. 906F1R_2
58. 940F1R
59. 951R
60. 951F1R
61. 978R
62. 978F1R
63. 978F1R_2
64. 1042R
Translation
